# Leaf pigments and photosystems stoichiometry underpin photosynthetic efficiency of related C_3_, C_3_-C_4_ and C_4_ grasses under shade

**DOI:** 10.1101/2020.10.08.330837

**Authors:** JV Sagun, WS Chow, O Ghannoum

**Affiliations:** ARC Centre of Excellence for Translational Photosynthesis, Hawkesbury Institute for the Environment, Western Sydney University, Hawkesbury Campus, Locked Bag 1797, Penrith, NSW 2751, Australia; ARC Centre of Excellence for Translational Photosynthesis, Research School of Biology, Australian National University, Canberra, ACT, 2601

**Author notes:** **Author contributions:** All authors conceived the research plans; JVS performed the experiments under the supervision of OG and WSC; JVS wrote the article with contribution from other authors.

**Keywords:** C_3_, C_3_-C_4_ and C_4_ grasses, C_4_ subtypes, chlorophyll fluorescence, leaf absorptance, leaf pigments, photosynthesis, quantum yield

## Abstract

The quantum yield of CO_2_ assimilation (QY) is generally lower in C_3_ relative to C_4_ plants at warm temperatures, and differs among the C_4_ subtypes. Here, we investigated whether variations in QY are linked to light absorption or conversion efficiency. We grew six representative grasses with C_3_, C_3_-C_4_ and C_4_ photosynthesis under full (control) or 20% (shade) sunlight, and measured the *in vivo* activity and stoichiometry of PSI and PSII, leaf spectral properties and pigment contents, and photosynthetic enzyme activities. Overall, shade reduced leaf photosynthesis, absorptance, especially in the green region, as well as carotenoids/chlorophylls and chlorophyll a/b ratios in C_4_ more than non-C_4_ species. Amongst C_4_, NADP-ME species had the highest QY and cyclic electron flow (CEF), and the NAD-ME species underwent the greatest reduction in leaf absorptance and pigments and PSII contents under shade, whist CEF and PSII/PSI were unaffected. These results demonstrate that the greater efficiency of the CO_2_ concentrating mechanism in NADP-ME grasses at low light depends on light absorption and harvesting properties in addition to coordination between the C_3_ and C_4_ cycles. This is important for maximising light energy absorption and providing the right ATP/NADPH ratio while minimising photoinhibition under variable light conditions.

**SUMMARY STATEMENT:** Changes in leaf absorptance, pigment contents and photosystems stoichiometry underpin photosynthetic efficiency and responses of closely related C_3_, C_3_-C_4_ and C_4_ grasses under shade

## INTRODUCTION

The main physiological trait predicting the dependence of C_3_ and C_4_ grass distribution on temperature is photosynthetic quantum yield, QY (Collatz *et al.*, 1998; Ehleringer, 1978; Ehleringer *et al.*, 1997). QY is the ratio of moles of CO_2_ assimilated to moles of quanta of photosynthetic active radiation (PAR) absorbed by a leaf (Ehleringer & Björkman, 1977). In normal air and warm temperatures (>30°C), QY is generally higher in C_4_ compared to C_3_ plants (Ehleringer & Björkman, 1977; Ludlow & Wilson, 1971; Maroco *et al.*, 1997) due to the higher energy cost of photorespiration relative to the cost of operating the CO_2_ concentrating mechanism (CCM) in C_4_ plants (Hatch, 1987). C_3_-C_4_ intermediate species have lower CO_2_ compensation points relative to C_3_ species due to a weak photorespiratory pump related to the localisation of the glycine decarboxylase activity in the bundle sheath cells (Rawsthorne, 1992; Sage, 2005). Consequently, these C_3_-C_4_ species tend to have intermediate QY relative to C_3_ and C_4_ species (Pinto *et al.*, 2011). In addition, QY differs among the C_4_ subtypes, with NADP-ME grasses generally having a higher QY, followed by PCK then NAD-ME grasses (Ehleringer & Pearcy, 1983; Sonawane *et al.*, 2018). In this study, we investigated whether QY variations are related to leaf absorptance and photosystems activity, especially under low irradiance.

QY partly depends on the stoichiometry of photosystem I (PSI), photosystem II (PSII) and associated light-harvesting and electron transport components in the chloroplasts (Anderson *et al.*, 1988; Chow *et al.*, 2012; Ghannoum *et al.*, 2005; Pfundel *et al.*, 1996; Romanowska *et al.*, 2008; Romanowska & Drożak, 2006). The PSII/PSI ratios differ between mesophyll cells (MC) and bundle sheath cells (BSC) of the C_4_ subtypes (Ghannoum *et al.*, 2005; Hatch, 1987; Hernández-Prieto *et al.*, 2019; Romanowska & Drożak, 2006). Several studies showed the dependence of QY on the PSII/PSI ratio in C_3_ plants (Chow *et al.*, 1990; Evans, 1987, 2006a) and in isolated MC and BSC chloroplasts from the three C_4_ subtypes (Romanowska & Drożak, 2006). These studies indicated that QY variation reflects the distribution of quanta between PSI and PSII, and the balance between these two photosystems in chloroplasts is a compensation strategy designed to correct unbalanced absorption of light. Such adjustments allow the plant to maintain high light use efficiency under diverse light conditions and confer a significant evolutionary advantage over that of a fixed photosystem stoichiometry in thylakoid membranes.

Light intensity during plant development is a key factor responsible for adjustments of the photosystems activity and stoichiometry (Anderson, 1986). In response to long-term shading, the photosynthetic apparatus acclimates to maximize light use efficiency (Boardman, 2003; Sage & McKown, 2006). Several factors constrain C_4_ photosynthesis under shade relative to C_3_ species. C_4_ plants may have lower QY under shade, especially at cooler temperatures, due to the additional energy requirement of the C_4_ pump (Ehleringer & Björkman, 1977; Krall & Pearcy, 1993). In particular, the CCM is less effective at low light due to increased relative photorespiration and BS CO_2_ leakiness which increase additional ATP demand (Bellasio & Griffiths, 2014; Henderson *et al.*, 1992; Kromdijk *et al.*, 2014; Kromdijk *et al.*, 2010; Kubásek *et al.*, 2007; Tazoe, 2008). The C_4_ photosynthetic machinery also lacks the capacity for maintaining a high state of photosynthetic induction during low-light periods (Horton & Neufeld, 1998; Sage & McKown, 2006).

In addition, the three C_4_ subtypes acclimate differently under shade. For example, shade (16% of full sunlight) compromised the CCM efficiency to a greater extent in NAD-ME than in PCK or NADP-ME C_4_ grasses by virtue of a greater increase in carbon isotope discrimination and a greater reduction in QY (Sonawane *et al.*, 2018). Together with other studies, it can be concluded that QY and CCM efficiency of NADP-ME species were not adversely affected by low light (Bellasio & Griffiths, 2014a; Bellasio & Lundgren, 2016; Ghannoum *et al.*, 2005; Yin & Struik, 2015). This was supported by a modelling approach which showed that the NADP-ME biochemical pathway is favoured at low light (Wang *et al.*, 2014). However, it is still unclear how these observed differences under shade relate to the changes in light reactions especially among C_4_ subtypes. Irradiance is likely to induce changes in the light reaction components and pigmentation which can alter relative light absorption and energy conversion efficiency by PSI and PSII. These changes might differ among photosynthetic types or C_4_ subtypes.

In this study, six representative species of C_3_, C_3_-C_4_ and the three subtypes of C_4_ photosynthesis were grown under high-light (control) and shade (20% sunlight) to investigate differences in plasticity of the light reactions to long-term exposure to shade. We asked if the high photosynthetic QY observed in C_4_ grasses and the variations in QY among the subtypes are associated with differences in the activity of the photosystems or pigment composition. In particular, we measured the stoichiometry of the two photosystems and relative electron fluxes *in vivo* in order to reflect functionality *in situ,* without any potential complication associated with isolation of thylakoids. For this purpose, we used both commercial equipment (Dual-PAM) which can simultaneously assess P700 and chlorophyll fluorescence, and custom–built equipment developed by (Chow *et al.*, 1989; Chow & Hope, 2004; Kou, *et al.*, 2013) which can measure the total electron flux through PSI (ETR1), the linear electron flux through PSII and PSI (LEF_O2_) in series, and cyclic electron flow around PSI (CEF). We then compared the results obtained using the commercial and custom-built systems. Finally, we measured activities of Rubisco and other enzymes involved in C_4_ photosynthesis to relate the observed QY to CCM efficiency under variable growth light conditions.

## MATERIALS AND METHODS

### Plant culture

Representative species of Paniceae grasses belonging to C_3_ (*Panicum bisulcatum*); C_3_-C_4_ intermediate (*Panicum milioides*); and C_4_ photosynthesis with NADP-ME subtype (*Panicum antidotale* and *Zea mays*); NAD-ME subtype (*Panicum miliaceum*); and PCK subtype (*Megathyrsus maximus*) were grown in vermiculite in a naturally lit greenhouse chamber at the Australian National University as described in (Sagun *et al.*, 2019). The chamber temperature was maintained at 24/21°C for day/night by an in-built greenhouse temperature control system. Within the chamber, a steel structure was erected and covered with shade cloth. The average ambient photosynthetic photon flux density, PPFD and temperature during the mid-day were 800 and 200 μmol photons m^−2^ s^−1^ and 30 and 29°C for the sun/control and shade treatments, respectively. Plants were watered regularly and fertilized with Osmocote® (Scotts Australia). Leaves were analysed from 4–5 week-old plants.

### Leaf gas exchange and chlorophyll fluorescence measurements

Leaf gas exchange was measured using an open gas exchange system (LI-6400; LICOR, Inc., Lincoln, NE, USA). Measurements were made at PPFD of 1000 (high-light or HL) and 200 μmol photons m^−2^ s^−1^ (shade or LL) and 28°C leaf temperature about 7-8 weeks after transplanting on an attached, youngest fully expanded leaf on the main stem. CO_2_ concentration were maintained at 400 μl L^−1^ within the leaf chamber, corresponding to the ambient level.

### Electrochromic signal to determine the photosystem stoichiometry

The electrochromic shift (ECS) signal from leaf segments (30 mm x 30 mm) was measured as described by (Chow & Hope, 2004). A measuring beam at 520 nm, selected by an interference filter (full width at half peak height = 2 nm), was admitted by an electronic shutter shortly before each measurement. The measuring beam was transmitted to the end window of a photomultiplier, through a leaf segment oriented at 45° to the light path. Shortly afterwards, data acquisition commenced before the triggering of a xenon single-turnover flash. The xenon flash was directed at 45° to the leaf segment, but at 90° to the measuring beam. Data acquisition continued for 50 ms in total, followed by closing of the electronic shutter to keep the leaf segment in darkness until the next measurement. Timing of these events was controlled by a pulse/delay generator (Model 555, Berkeley Nucleonics Corporation, USA). The xenon flash was filtered by a red Perspex long-pass filter (transmitted wavelengths >590 nm) and a heat-reflection filter that did not transmit wavelengths >700 nm. The elimination of the far-red component of the xenon flash by the heat-reflection filter was meant to avoid unbalanced excitation of both photosystems, since a far-red component, if present, would reach further into the leaf tissue where it would excite predominantly PSI. It is desirable to excite both photosystems in the same tissue, particularly if the flash is only sub-saturating. Xenon flashes were given at 0.2 Hz; typically, 25 signals were averaged to improve the signal-to-noise ratio.

The ECS signal was also measured in leaf segments during suppression of PSI by moderately strong far-red light (Figure S1). Leaf segments were illuminated for 10 s to steady-state by an array of far-red, light-emitting diodes in conjunction with a Schott RG9 filter (~695 μmol photons m^−2^ s^−1^, peak wavelength 741 nm, range 700–780 nm). The direction of the moderately strong far-red light made an angle of ~45° with the leaf segment. The P700 molecules in PSI reaction centres were not all photo-oxidized, however, necessitating correction for a small fraction of P700 in the reduced state. The small fraction of reduced P700 was determined from the redox state of P700 during illumination with the same far-red light as described by (Chow & Hope, 2004) (Figure S2).

The complementary fraction *r* of reduced P700 was then obtained and taken into account when evaluating the contribution of PSII reaction centres to the ECS signal in the presence of moderately strong far-red light. Let the contents of PSII and PSI reaction centres be *n*_PSII_ and *n*_PSI_, respectively. The amplitude of the fast ECS rise, either in the absence (*E*_f(−FR)_) or the presence (*E*_f(+FR)_) of far-red light is directly proportional to the sum of the contributing reaction centres, with *k* as the inverse of the constant of proportionality:

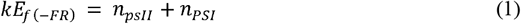

and

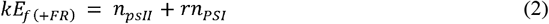

Manipulation of these two equations gives the stoichiometry of the two photosystems:

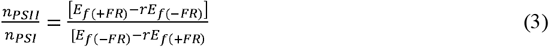

### Quantification of functional PSII by the oxygen yield per single turn-over flash

The same leaf disc was used for quantification of the functional PSII centres, by measuring the amount of oxygen evolution per single turn-over flash in a leaf-disc oxygen electrode, with continuous background far-red light (Chow *et al.*, 1989). These estimations are based on the assumption that for every four flashes (at 10 Hz), four electrons are transferred through each functional PSII, resulting in the evolution of one O_2_ molecule.

### Estimation of steady-state cyclic electron flux (CEF) around PSI using custom-built equipment

The rate of CEF was estimated using the following equation:

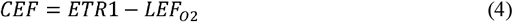

where ETR1 is the total electron flux through PSI and LEF_O2_ is linear electron flux through both photosystems measured as the gross rate of O_2_ evolution. Eqt 4 gives an estimate of CEF because ETR1 could include other electron fluxes going through PSI if they exist (e.g., charge separation followed by charge recombination to the ground state – a kind of local cycle within PSI. LEF_O2_ and ETR1 were measured as described below:

### LEF_O2_ measurement

LEF_O2_ was estimated by measuring O_2_ evolution in a gas-phase oxygen electrode (Hansatech, King’s Lynn, UK) chamber, thermostat-controlled at 25°C and with a multifurcated light guide, containing 1% CO_2_ supplied by fabric matting moistened with 1 M NaHCO_3_/Na_2_CO_3_ (pH 9). White incandescent light from a projector halogen lamp, filtered by a Calflex C heat-reflecting filter (Linos Photonics, Göttingen, Germany) and neutral-density filters, was used to illuminate a leaf disk. O_2_ evolution was measured over several minutes until it reached a steady state. The post-illumination drift was subtracted algebraically from the steady-state net oxygen evolution rate, and the gross O_2_ evolution rate obtained was multiplied by four to give LEF_O2_. For calibration of the oxygen signals, 1 mL of air at 25°C (taken to contain 8.05 μmol O_2_) was injected into the gas-phase O_2_ electrode chamber.

### ETR1 measurement from redox kinetics of P700

Redox changes of P700 were observed with a dual wavelength (820/870 nm) unit (ED-P700DW) attached to a pulse amplitude modulation (PAM 101) fluorometer (Walz, Effeltrich, Germany) in the reflectance mode (response time constant = 95 μs) as described by (Kou *et al.*, 2013). A leaf disc was brought to steady-state photosynthesis by illuminating it with white actinic light for about 10 min before simultaneous measurement of O_2_ evolution and chlorophyll fluorescence yields. To retain steady-state for P700^+^ measurements, immediately after these simultaneous measurements, each leaf disc was re-illuminated with the same actinic light for 9.016 s, using an electronic shutter controlled by one terminal of a pulse/delay generator (Model 555, Berkeley Nucleonics, San Rafael, CA, USA).

During each 9.016-s illumination, when the photo-oxidized P700 signal reached a steady level *P*, at time T= 8.80 s (corresponding to the time point t = −50 ms), data acquisition (using software written by the late AB Hope) was started by a second terminal of the pulse/delay generator. At T= 8.85 s, a strong far-red light (FR, ~2000 μmol m^−2^ s^−1^) from a light-emitting diode array (741 nm ± 13 nm, LED735–66–60, Roithner LaserTechnik, Vienna, Austria) was triggered on for 100 ms using a third terminal of the pulse/delay generator. The strong FR light depleted electrons from the inter-system chain, so that the subsequent saturating pulse maximally oxidised P700 (Siebke *et al.*, 1997). While the strong FR light was on, at T= 8.90 s, a saturating light pulse (~9000 μmol m^−2^ s^−1^) was applied for 10 ms by a pulse from a fourth terminal of the pulse/ delay generator, yielding the maximally-oxidised 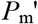 signal (where 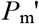 is the maximum P700^+^ signal in actinic light). Finally, the white actinic light was turned off by the electronic shutter (at T= 9.016 s). Data acquisition continued for 85 ms after cessation of actinic illumination to obtain the baseline corresponding to complete reduction of P700. Immediately after completion of one cycle of illumination and data acquisition (at T= 9.101 s), another 9.016-s illumination was restarted, thereby maintaining steady-state photosynthesis. Nine traces were averaged automatically to improve the signal-to-noise ratio. Next, the maximum photo-oxidisable P700 content in the absence of actinic light, *P*_m_, was determined (Figure S2). A steady-state attained by illumination with weak continuous far-red light (~12 μmol m^−2^ s^−1^, 723 nm) for >10 s was then established. A single saturating turnover flash was then superimposed. Flashes were given at 0.2 Hz; nine consecutive signals were automatically averaged. The maximum P700^+^ signal (*P*_m_) immediately after the flash was used to normalise the signal interval 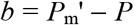. The photochemical yield of PSI [Y(I)] is then given by:

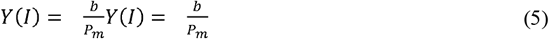

where *P*_m_ is the maximum signal, and *b* is the signal interval (Klughammer & Schreiber, 2008).

ETR1 was then calculated as:

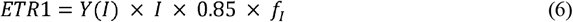

where *I* is the irradiance, 0.85 is the assumed absorptance and *f*_I_ is the fraction of absorbed white light partitioned to PSI which was experimentally derived (Sagun *et al.*, 2019).

### Measurements of leaf reflectance, transmittance and absorptance

Leaf reflectance, transmittance and absorptance to solar radiation over the 400–700 nm wavelength were measured with a Li-Cor 1800-12 Integrating Sphere (Li-Cor, Inc., Lincoln, NE, USA), coupled by a 200 μm diameter single mode fibre to an Ocean Optics model USB2000 spectrometer (Ocean Optics Inc., Dunedin, FL, USA), with a 2048 element detector array, 0.5 nm sampling interval, and 7.3 nm spectral resolution in the 350–1000 nm range. The Ocean Optics software is designed for signal verification, adjustment of integration time, and data acquisition. An integration time of 13 ms was used for all sample measurements. Single leaf reflectance and transmittance measurements were acquired following the methodology described in the product manual for the LiCor 1800-12 system (Li-Cor Inc., 1984). Reflectance of the sample (*R*_s_) is the amount of flux reflected by the leaf, normalized by the amount of flux incident on it and calculated as:

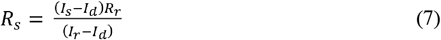

where *I*_s_ is the measured sphere output when the sample is illuminated, *I*_r_ is that measured when the reference material (barium sulfate) is illuminated and Id is measured by illuminating the sample port with no sample in place. For equation (6), it was assumed that the reflectance of the reference material (*R_r_*) is 1. Transmittance of the sample (*T_s_*) is the amount of flux transmitted by the leaf, normalized by the amount of flux incident on it and calculated as:

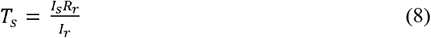

Any flux not reflected or transmitted is absorbed (*A_s_*), and the absorptance was obtained using the following equation based on conservation of energy:

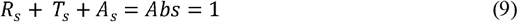

### Chlorophyll fluorescence using Dual-PAM

The chlorophyll (Chl) fluorescence and P700 redox state measurement were also determined *in vivo* with the same leaves using a Dual-PAM-100 (Heinz Walz). The *F_v_/F_m_* and *P_m_* were determined after dark adaptation for 20 minutes. Light responses of leaf chlorophyll fluorescence and P700 were also measured after 15 min light adaptation under PPFD of 1000 μmol photons m^−2^ s^−1^. Light-adapted fluorescence parameters were recorded after 3 min exposure to each of the PPFDs (30, 37, 46, 61, 77, 94, 119, 150, 190, 240, 297, 363, 454, 555, 684, 849, 1052, 1311, 1618 and 1976 μmol photons m^−2^ s^−1^).

The fluorescence parameters were calculated as follows:

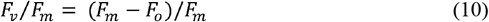

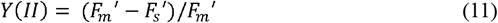

where *F*_o_ represents the minimum fluorescence in the dark-adapted state and light-adapted state, respectively, and *F*_m_ and 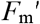 represent the maximum fluorescence in the dark-adapted state and actinic-light-adapted state measured upon illumination with a pulse (600 ms) of saturating light (10000 μmol photons m^−2^ s^−1^), respectively (Kramer *et al.*, 2004). 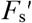 is the light steady-state fluorescence. *F*_v_/*F*_m_ represents the maximum quantum yield of PSII in the dark-adapted state. Y(II) is the effective quantum yield of PSII.

The parameters related to PSI are calculated as follows:

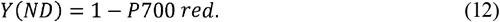

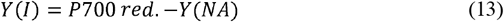

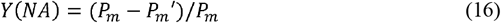

where *P*700 red. is the fraction of total P700 in the reduced state. After far-red pre-illumination for 10 s, *P*_m_ was determined through the application of a saturation pulse. 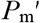 was determined similarly to *P*_m_, in actinic light but without far-red pre-illumination. This method was taken from Klughammer & Schreiber (2008). Y(ND) represents the fraction of total P700 that is oxidized in a given state due to a lack of donors which is enhanced by a trans-thylakoid proton gradient (photosynthetic control at the cytochrome *b_6_f* complex as well as down-regulation of PSII) and photo-damage to PSII. Y(NA) represents the fraction of overall P700 that cannot be oxidized by a saturation pulse in a given state due to a lack of acceptors; it is enhanced by dark adaptation (deactivation of key enzymes of the Calvin–Benson cycle) and damage at the site of CO_2_ fixation. The saturating pulse used for P700 measurements was 10000 μmol m^−2^ s^−1^. CEF was also calculated using equation (4).

### Quantum yield for CO_2_ uptake and assimilation quotient calculations

Apparent quantum yield for CO_2_ uptake was calculated as the ratio of CO_2_ assimilation rates measured at HL (*QY*_1000_) or at growth irradiance (*QY*_growth_) to absorbed irradiance as follows:

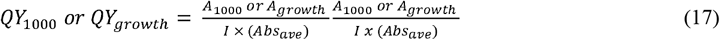

where *A*_1000_ and *A*_growth_ are the CO_2_ assimilation rates at 1000 μmol photons m^−2^ s^−1^ and at growth irradiance, *I* is the measurement irradiance, and *Abs_ave_* is the average leaf absorptance for each species and light treatment in the blue (470 nm) and red (660 nm) wavelength regions (Figures 3A-F), which correspond to the LED light source of the Li-Cor equipment.

The assimilation quotient (*AQ*_1000_) was calculated using CO_2_ assimilation (measured at 1000 photons μmol m^−2^ s^−1^ and ambient CO_2_ 400 ppm) and O_2_ evolution rates (measured at saturating light of 2000 μmol m^−2^ s^−1^ and saturating CO_2_ of >1000 ppm). Given *A* and LEF_O2_ were not measured under the same conditions, these values should be considered with caution.

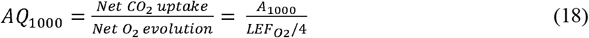

where LEF_O2_ has units μmol electrons m^−2^ s^−1^.

### Carotenoid and chlorophyll analysis and quantification by HPLC

Carotenoids and chlorophyll analysis through HPLC was done following the modified protocol from Pogson *et al.* (1996). Approximately 50 mg fresh leaves were extracted in a microcentrifuge tube by grinding using a tissue lyser (Qiagen) (25 Hz for 3 minutes). Pigments were extracted by adding 1 mL of extraction buffer, comprising of acetone:ethyl acetate (60:40) with 0.1% Butylated hydroxytoluene (BHT) and then vortexed for 1 minute or until the tissue lost its green colour. The organic component of the extract was separated by gently adding 900 μL of water and inverting the tube 4 – 5 times. The solution was then centrifuged for 5 minutes at 15000 rpm at 4°C. The upper phase was collected and transferred in a new microcentrifuge tube (a little of the lower phase was still obvious at this point). This extract was then centrifuged for 5 minutes at 15000 rpm at 4°C and 200 μL was transferred to another tube. A final volume of 20 μL was used for injection into HPLC system (Agilent 1260 Infinity) equipped with YMC-C30 (250 × 4.6 mm, S-5μm) column and Diode Array Detector (DAD) detector. A newly optimized 17 minutes elution method was used to separate pigments using reverse phase solvent gradient comprised of mobile phase A (methanol:water:triethylamine, 98:2:0.1 v/v) and B (methyl tert-butyl ether). The initial solvent composition was 85% and 15% of the solvent A and B, respectively, for one minute, followed by 35% of B up to minute 11. The ratio of B increased to 65% at 11.1 minutes followed by a gradient up to 70% at 15 minutes and stabilized at this ratio through to 17 minutes. The solvent flow rate was 1 ml/min until 17 minutes from the beginning. Solvent composition was returned to the initial state (15% of B) at 17.1 minutes and the column was equilibrated up to 25 minutes with the flow rate of 2.0 ml/min. The column temperature was maintained at 23°C and auto-sampler temperature was set to 8°C and illumination turned off. Carotenoid and chlorophyll peak signals were analysed at 440 nm based on Pogson *et al.* (1996) and Cuttriss *et al.* (2007). Figure S3 shows the typical HPLC profiles of leaf pigments absorbing at 440 nm, with peaks corresponding to different carotenoids and chlorophylls.

### Activity of Rubisco, PEPC, NADP-ME, NAD-ME and PCK

Following gas exchange measurements, leaf discs (0.6 cm^2^) were taken and rapidly frozen in liquid nitrogen. Activities of photosynthetic enzymes were measured at 25°C using an NADH-coupled enzyme assay with the rate of NADH oxidation or reduction monitored at 340 nm using a diode array spectrophotometer (Agilent model 8453) as described by Sharwood *et al.* (2016). For assays of Rubisco, PEPC and NADP-ME, leaf discs were extracted using ice-cold mortar and pestle into 1 ml of ice cold, extraction buffer [50 mM EPPS-NaOH, pH 7.8, 1 mM EDTA, 5 mM DTT, 5 mM MgCl_2_, 1% (v/v) plant protease inhibitor cocktail (Sigma-Aldrich), and 1% (w/v) polyvinylpolypyrrolidone (PVPP). The extract was rapidly centrifuged for 30 seconds at 15,000 × *g* at 4°C. Protein content was measured against BSA standards using Pierce Coomasie Plus (Bradford) protein assay kit (Thermoscientific, Rockford, USA). Fifty μL of the soluble leaf protein was used to measure Rubisco content by [^14^C]CABP (2-C-carboxyarabinitol 1,5-bisphosphate) binding as described by Sharwood *et al.*, (2016). Rubisco activity was measured in assay buffer [100 mM EPPS-NaOH, pH 8.0, 10 mM MgCl_2_, 0.2 mM NADH, 20 mM NaHCO_3_, 1 mM ATP (pH 7.0), 5 mM phosphocreatine (pH 7.0), 50 U creatine phosphokinase, 0.2 mg carbonic anhydrase, 50 U 3-phosphoglycerate kinase, 40 U glyceraldehyde-3-phosphate dehydrogenase, 113 U triose-phosphate isomerase, 39 U glycerol-3-phosphate dehydrogenase]. Ribulose-1, 5-bisphosphate (RuBP) (0.26 mM) was included in the cuvettes, with 10 μl of soluble leaf protein extract added to start the assays. Maximal PEPC activity was measured in assay buffer containing [50 mM EPPS-NaOH (pH 8.0), 0.5 mM EDTA, 10 mM MgCl_2_, 0.2 mM NADH, 5 mM glucose-6-phosphate, 1 mM NaHCO_3_, 1 U/mL MDH] after the addition of 8 mM of PEP. NADP-ME activity was measured in assay buffer containing [50 mM Tricine buffer (pH 8.3), 0.5 mM NADP, 5 mM malate, 0.1 mM EDTA] after addition of 10 mM MgCl_2_.

The maximal activity of PCK was measured in the carboxylase direction using a method described by Sharwood *et al.* (2016) in an NADH-coupled assay. Leaf discs (0.6 cm^2^) were extracted in 50 mM HEPES pH 7.0, 5 mM DTT, 1% (w/v) PVPP, 2 mM EDTA, 2 mM MnCl_2_, and 0.05% Triton using ice-cold mortar and pestle. PCK activity from leaf extracts was measured in assay buffer [50 mM HEPES, pH 7.0, 4% mercaptoethanol (w/v), 100 mM KCl, 90 mM NaHCO_3_, 1 mM ADP, 2 mM MnCl_2_, 0.14 mM NADH, and malate dehydrogenase (MDH; 6 U; 3.7 μL)] after the addition of 15 mM PEP. The activity of NAD-ME was measured using a method described by Setién *et al.* (2014). Leaf discs (0.6 cm^2^) were extracted in [50 mM HEPES-KOH (pH 8.0), 2 mM EDTA, 0.05% Triton, 2 mM MnCl_2_, 10 mM DTT, 1% PVPP and 1% (v/v) plant protease inhibitor cocktail in ice-cold mortar and pestle. NAD-ME activity was measured at 25°C in a reaction buffer containing [50 mM HEPES-KOH (pH 8.0), 5 mM NAD, 5 mM DTT, 0.1 mM coenzyme-A, 5 mM malate, 0.2 mM EDTA] after the addition of 4 mM MnCl_2_.

### Data analysis

Leaf parameters were measured on area, Chl, and/or Rubisco basis. For each variable, four replicates (independent samples) were obtained for the two light treatments. The results were subjected to analysis of variance and the means were compared by the Tukey test at 5% probability.

## RESULTS

### Photosynthetic rate and functional PSII content

The control C_4_ species generally had higher CO_2_ assimilation rate measured at their growth light (HL, 1000 μmol photons m^−2^ s^−1^) (*A*_growth_) followed by the C_3_-C_3_ then C_3_ species (Figure 1A; Tables 1 and 2). Shade reduced *A*_growth_ measured at growth light (LL, 200 μmol photons m^−2^ s^−1^) to a greater extent in the C_4_ and C_3_-C_3_ (−50% to −60%) relative to the C_3_ species (−20%). Yet, *A*_growth_ remained slightly higher in the C_4_ relative to non-C_4_ species under (Figure 1A; Tables 1 and 2). Under control conditions, the C_4_ species had lower amount of functional PSII in the leaf relative to the C_3_ and C_3_-C_3_ species (Figure 1B; Tables 1 and 2). Shade reduced the amount of leaf PSII in all species; and this reduction was largest in *P. miliaceum* (NAD-ME) such that it had the lowest amount under shade (Figure 1B; Tables 1 and 2).

**Figure 1.**
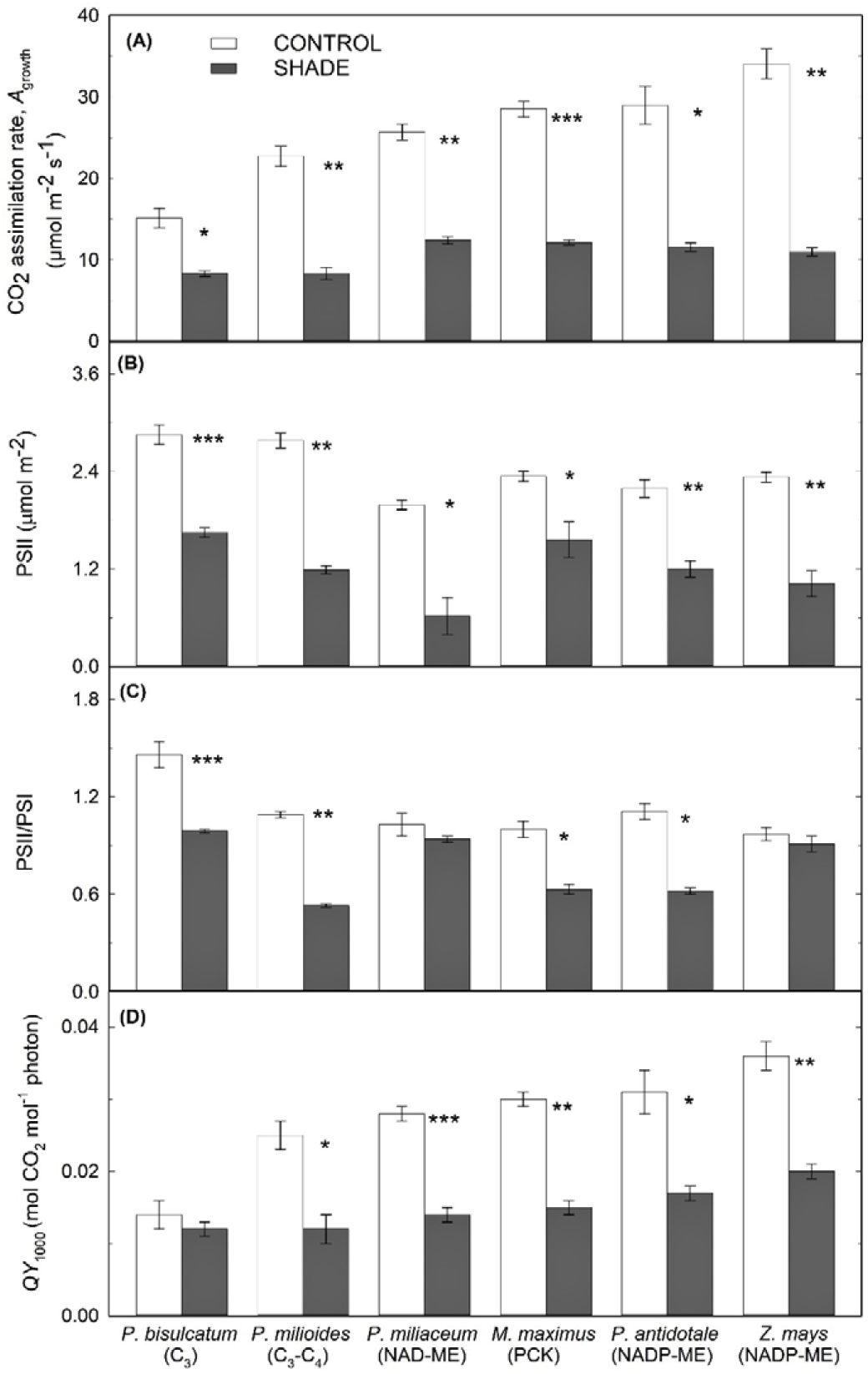
CO_2_ assimilation, functional PSII content, PSII/PSI ratio, and the quantum yield for CO_2_ uptake. Leaf (**A**) CO_2_ assimilation rate (*A*growth) measured at ambient CO_2_ and growth irradiance (1000 or 200 μmol photons m^−2^ s^−1^ for HL and LL plants, respectively); (**B**) functional PSII content (determined by the oxygen yield per single turnover); **(C)** PSII/PSI ratio (determined by electrochromic shift); and **(D)** photosynthetic quantum yield at HL (*QY*1000) for control and shade-grown C_3_, C_3_-C_4_ and C_4_ grasses. Statistical significance levels (t-test) for the growth condition within each species are shown and they are: * ≡ *p*<0.05; ** ≡ *p* < 0.01; *** ≡ *p* < 0.001. Each column represents the mean ± s.e. of each species (*n* = 4 plants).

**Table 1.** Statistical summary. Summary of statistical analysis using two-way ANOVA for the effects of shade on the leaf parameters of six grass species grown under full sunlight (~800 μmol photons m^−2^ s^−1^) or shaded (~200 μmol photons m^−2^ s^−1^) conditions. See Tables and Figures for definition of the various parameters.

| Parameter | Main effects (P) |  | Interactions (P) |
| --- | --- | --- | --- |
|  | Species | Treatment | Species x Treatment |
| PSII content ($\mu\text{mol m}^{-2}$) | 0.000 | 0.000 | 0.037 |
| PSI content ($\mu\text{mol m}^{-2}$) | 0.000 | 0.000 | 0.000 |
| PSII/PSI ratio | 0.000 | 0.000 | 0.000 |
| Absorptance at 470 nm | 0.000 | 0.000 | 0.046 |
| Absorptance at 550 nm | 0.000 | 0.000 | 0.026 |
| Absorptance at 665 nm | 0.000 | 0.000 | 0.004 |
| Neoxanthin (%) | 0.000 | 0.020 | 0.249 |
| Violaxanthin (%) | 0.000 | 0.000 | 0.000 |
| Antheraxanthin (%) | 0.046 | 0.341 | 0.176 |
| Lutein (%) | 0.000 | 0.000 | 0.342 |
| Zeaxanthin (%) | 0.000 | 0.132 | 0.000 |
| β-carotene (%) | 0.000 | 0.000 | 0.049 |
| Chlorophyll *a/b* | 0.000 | 0.000 | 0.580 |
| Total carotenoids ($\text{mg m}^{-2}$) | 0.004 | 0.000 | 0.000 |
| Total chlorophyll ($\text{mg m}^{-2}$) | 0.000 | 0.000 | 0.000 |
| Total carotenoids/total chlorophyll | 0.000 | 0.000 | 0.000 |
| $F_v/F_m$ | 0.000 | 0.000 | 0.017 |
| Dual-PAM Y(I) ($\mu\text{mol e}^{-} \text{m}^{-2} \text{s}^{-1}$) | 0.000 | 0.000 | 0.002 |
| Dual-PAM Y(II) ($\mu\text{mol e}^{-} \text{m}^{-2} \text{s}^{-1}$) | 0.000 | 0.000 | 0.002 |
| Dual-PAM ETR1 ($\mu\text{mol e}^{-} \text{m}^{-2} \text{s}^{-1}$) | 0.003 | 0.000 | 0.002 |
| Dual- PAM ETR2 ($\mu\text{mol e}^{-} \text{m}^{-2} \text{s}^{-1}$) | 0.000 | 0.000 | 0.003 |
| Dual-PAM CEF ($\mu\text{mol e}^{-} \text{m}^{-2} \text{s}^{-1}$) | 0.000 | 0.001 | 0.023 |
| Y(I) | 0.000 | 0.000 | 0.000 |
| ETR1 ($\mu\text{mol e}^{-} \text{m}^{-2} \text{s}^{-1}$) | 0.000 | 0.000 | 0.000 |
| LEF<sub>O2</sub> ($\mu\text{mol e}^{-} \text{m}^{-2} \text{s}^{-1}$) | 0.000 | 0.000 | 0.000 |
| CEF ($\mu\text{mol e}^{-} \text{m}^{-2} \text{s}^{-1}$) | 0.000 | 0.000 | 0.000 |
| CEF/ETR1 | 0.000 | 0.000 | 0.000 |
| $A_{1000}$ ($\mu\text{mol CO}_2 \text{m}^{-2} \text{s}^{-1}$) | 0.000 | 0.000 | 0.001 |
| $A_{\text{growth}}$ ($\mu\text{mol CO}_2 \text{m}^{-2} \text{s}^{-1}$) | 0.000 | 0.000 | 0.000 |
| $AQ_{1000}$ ($\text{mol CO}_2 \text{mol}^{-1} \text{O}_2$) | 0.000 | 0.088 | 0.216 |
| $QY_{1000}$ ($\text{mol CO}_2 \text{mol}^{-1} \text{photons}$) | 0.000 | 0.000 | 0.000 |
| $QY_{\text{growth}}$ ($\text{mol CO}_2 \text{mol}^{-1} \text{photons}$) | 0.000 | 0.000 | 0.000 |
| Rubisco activity ($\mu\text{mol m}^{-2} \text{s}^{-1}$) | 0.000 | 0.000 | 0.000 |
| $K_{\text{cat}}$ ($\text{s}^{-1}$) | 0.000 | 0.000 | 0.025 |
| Rubisco content ($\mu\text{mol sites m}^{-2}$) | 0.000 | 0.000 | 0.000 |
| Protein content ($\text{g m}^{-2}$) | 0.001 | 0.009 | 0.167 |
| PEPC activity ($\mu\text{mol m}^{-2} \text{s}^{-1}$) | 0.000 | 0.000 | 0.660 |
| PEPC activity/Rubisco activity | 0.000 | 0.000 | 0.000 |
| NADP-ME activity ($\mu\text{mol m}^{-2} \text{s}^{-1}$) | 0.000 | 0.000 | 0.000 |
| PCK activity ($\mu\text{mol m}^{-2} \text{s}^{-1}$) | 0.000 | 0.000 | 0.001 |
| NAD-ME activity ($\mu\text{mol m}^{-2} \text{s}^{-1}$) | 0.000 | 0.000 | 0.000 |
| Total decarboxylases ($\mu\text{mol m}^{-2} \text{s}^{-1}$) | 0.000 | 0.000 | 0.000 |

**Table 2.** Summary of photosynthetic parameters in representative species of C_3_, C_3_-C_4_, and C_4_ grasses. Photosynthetic rates were measured at growth irradiance, *A*_growth_ (1000 or 200 μmol photons m^−2^ s^−1^) using Licor. Electron transport parameters were measured under saturating light (2000 μmol photons m^−2^ s^−1^) using custom-built equipment. Representative species of C_3_, C_3_-C_4_, and C_4_ grasses were grown under control (~800 μmol photons m^−2^ s^−1^) or shaded (~200 μmol photons m^−2^ s^−1^) conditions. Y(I) is the effective quantum yield of PSI, LEF_O2_ is the total electron flux, ETR1 is the electron transport rate through PSI, CEF is the cyclic electron flux around PSI, *AQ_1000_* is the assimilation quotient (ratio of A_1000_ to LEF_O2_), and *QY* is the ratio of *A* (*A*_growth_ or *A*_1000_) to absorbed light. Different letters indicate significantly different (*p*<0.05) ranking within each species using multiple-comparison Tukey’s post-hoc test. Values are means ± s.e. of species (*n* = 4 plants).

| Parameter | Light | *P. bisulcatum* | *P. milioides* | *P. miliaceum* | *M. maximus* | *P. antidotale* | *Z. mays* |
| --- | --- | --- | --- | --- | --- | --- | --- |
|  |  | C<sub>3</sub> | C<sub>3</sub>-C<sub>4</sub> | NAD-ME | PCK | NADP-ME | NADP-ME |
| $A_{\text{growth}}$<br>($\mu\text{mol m}^{-2} \text{s}^{-1}$) | CONTROL | 15.1 ± 1.2 d | 22.7 ± 1.2 c | 25.7 ± 1.0 bc | 28.5 ± 1.0 b | 29.01 ± 2.35 ab | 34.06 ± 1.86 a |
| | SHADE | 8.33 ± 0.33 ab | 8.3 ± 0.75 b | 12.42 ± 0.44 a | 12.13 ± 0.29 ab | 11.57 ± 0.52 ab | 10.99 ± 0.49 ab |
| | Shade/Control | 0.8 ± 0.13 a | 0.37 ± 0.07 b | 0.5 ± 0.02 b | 0.43 ± 0.02 b | 0.41 ± 0.06 b | 0.33 ± 0.04 b |
| PSII content<br>($\mu\text{mol m}^{-2}$) | CONTROL | 2.85 ± 0.12 a | 2.78 ± 0.09 a | 1.99 ± 0.06 b | 2.34 ± 0.06 ab | 2.19 ± 0.11 b | 2.33 ± 0.06 ab |
| | SHADE | 1.65 ± 0.06 a | 1.19 ± 0.05 ab | 0.62 ± 0.23 c | 1.56 ± 0.22 a | 1.20 ± 0.10 ab | 1.02 ± 0.16 bc |
| | Shade/Control | 0.58 ± 0.02 ab | 0.43 ± 0.03 ab | 0.31 ± 0.12 b | 0.67 ± 0.1 a | 0.55 ± 0.06 ab | 0.43 ± 0.06 ab |
| PSII/PSI | CONTROL | 1.46 ± 0.08 a | 1.09 ± 0.02 b | 1.03 ± 0.07 b | 1.00 ± 0.05 b | 1.11 ± 0.05 b | 0.97 ± 0.04 b |
| | SHADE | 0.99 ± 0.01 a | 0.53 ± 0.01 b | 0.94 ± 0.02 a | 0.63 ± 0.03 b | 0.62 ± 0.02 b | 0.91 ± 0.05 a |
| | Shade/Control | 0.68 ± 0.03 b | 0.48 ± 0.01 b | 0.93 ± 0.07 a | 0.64 ± 0.05 b | 0.56 ± 0.04 b | 0.94 ± 0.06 a |
| PSI content<br>($\mu\text{mol m}^{-2}$) | CONTROL | 1.97 ± 0.10 b | 2.55 ± 0.11 a | 1.96 ± 0.13 b | 2.35 ± 0.15 b | 1.99 ± 0.15 ab | 2.41 ± 0.11 ab |
| | SHADE | 1.67 ± 0.06 ab | 2.27 ± 0.11 a | 0.66 ± 0.25 c | 2.46 ± 0.33 a | 1.94 ± 0.18 ab | 1.14 ± 0.23 bc |
| | Shade/Control | 0.85 ± 0.03 abc | 0.90 ± 0.08 ab | 0.37 ± 0.17 c | 1.06 ± 0.17 a | 0.97 ± 0.05 a | 0.47 ± 0.08 bc |
| Y(I) | CONTROL | 0.18 ± 0 e | 0.24 ± 0.01 c | 0.21 ± 0 d | 0.15 ± 0 e | 0.31 ± 0.01 b | 0.37 ± 0.02 a |
| | SHADE | 0.08 ± 0 bc | 0.1 ± 0 ab | 0.09 ± 0.01 b | 0.05 ± 0.01 c | 0.07 ± 0 bc | 0.12 ± 0.01 a |
| | Shade/Control | 0.44 ± 0.02 a | 0.39 ± 0.01 ab | 0.41 ± 0.03 ab | 0.35 ± 0.04 ab | 0.22 ± 0.01 c | 0.33 ± 0.01 b |
| $\text{LEF}_{\text{O}_2}$<br>($\mu\text{mol e}^- \text{m}^{-2} \text{s}^{-1}$) | CONTROL | 79.9 ± 5.1 c | 124.3 ± 3 b | 129.6 ± 2.2 b | 67.6 ± 1.1 d | 68.1 ± 2.1 d | 142.5 ± 1.4 a |
| | SHADE | 48.7 ± 0.2 bc | 48.1 ± 3.7 bc | 54.4 ± 2.6 b | 31.1 ± 1.3 d | 41.8 ± 1.2 c | 66 ± 1 a |
| | Shade/Control | 0.62 ± 0.04 a | 0.39 ± 0.03 b | 0.42 ± 0.03 b | 0.46 ± 0.01 b | 0.61 ± 0.02 a | 0.46 ± 0.01 b |
| ETR1<br>($\mu\text{mol e}^- \text{m}^{-2} \text{s}^{-1}$) | CONTROL | 152.2 ± 3.4 d | 207.4 ± 8.7 c | 218.1 ± 2.3 c | 153 ± 3.1 d | 320.8 ± 7.4 b | 373.1 ± 16.9 a |
| | SHADE | 53.7 ± 3.1 c | 64.8 ± 1.8 bc | 89.2 ± 4.9 b | 53.6 ± 6.7 c | 69.6 ± 1.2 bc | 121 ± 4.9 a |
| | Shade/Control | 0.35 ± 0.01 a | 0.31 ± 0.01 ab | 0.41 ± 0.03 a | 0.35 ± 0.04 a | 0.22 ± 0.01 b | 0.33 ± 0.01 a |
| CEF<br>($\mu\text{mol e}^- \text{m}^{-2} \text{s}^{-1}$) | CONTROL | 72.3 ± 8.2 b | 83.1 ± 7.6 b | 88.5 ± 3.8 b | 85.4 ± 4.1 b | 252.7 ± 7.3 a | 230.6 ± 17.3 a |
| | SHADE | 4.9 ± 3 b | 16.7 ± 2.7 b | 34.9 ± 6.8 ab | 22.6 ± 7.7 b | 27.8 ± 1 ab | 55 ± 4.9 a |
| | Shade/Control | 0.06 ± 0.04 b | 0.21 ± 0.05 ab | 0.41 ± 0.1 a | 0.27 ± 0.09 ab | 0.11 ± 0.01 b | 0.24 ± 0.01 ab |
| CEF/ETR1 | CONTROL | $0.47 \pm 0.04$ cd | $0.40 \pm 0.02$ d | $0.41 \pm 0.01$ d | $0.56 \pm 0.02$ bc | $0.79 \pm 0.01$ a | $0.62 \pm 0.02$ b |
| | SHADE | $0.08 \pm 0.05$ b | $0.26 \pm 0.05$ ab | $0.38 \pm 0.06$ a | $0.39 \pm 0.09$ a | $0.40 \pm 0.01$ a | $0.45 \pm 0.02$ a |
| | *Shade/Control* | $0.15 \pm 0.09$ b | $0.67 \pm 0.14$ ab | $0.96 \pm 0.18$ a | $0.69 \pm 0.16$ ab | $0.51 \pm 0.01$ ab | $0.73 \pm 0.02$ a |
| $AQ_{1000}$ (mol CO<sub>2</sub><br>mol<sup>-1</sup> O<sub>2</sub>) | CONTROL | $0.19 \pm 0.01$ b | $0.18 \pm 0.01$ b | $0.2 \pm 0.01$ b | $0.42 \pm 0.01$ a | $0.43 \pm 0.04$ a | $0.24 \pm 0.01$ b |
| | SHADE | $0.23 \pm 0.02$ b | $0.22 \pm 0.02$ b | $0.21 \pm 0.01$ b | $0.47 \pm 0.03$ a | $0.38 \pm 0.03$ a | $0.28 \pm 0.01$ b |
| | *Shade/Control* | $1.05 \pm 0.1$ a | $1.35 \pm 0.13$ a | $1.23 \pm 0.08$ a | $1.24 \pm 0.1$ a | $0.86 \pm 0.1$ a | $1.04 \pm 0.07$ a |
| $QY_{1000}$ (mol CO<sub>2</sub><br>mol<sup>-1</sup> photons) | CONTROL | $0.014 \pm 0.002$ c | $0.025 \pm 0.002$ b | $0.028 \pm 0.001$ ab | $0.030 \pm 0.001$ ab | $0.031 \pm 0.003$ ab | $0.036 \pm 0.002$ a |
| | SHADE | $0.012 \pm 0.001$ b | $0.012 \pm 0.002$ ab | $0.014 \pm 0.001$ ab | $0.015 \pm 0.001$ ab | $0.017 \pm 0.001$ a | $0.020 \pm 0.001$ a |
| | *Shade/Control* | $0.89 \pm 0.11$ a | $0.51 \pm 0.1$ b | $0.49 \pm 0.02$ b | $0.52 \pm 0.03$ b | $0.57 \pm 0.06$ b | $0.57 \pm 0.04$ b |
| $QY_{\text{growth}}$ (mol CO<sub>2</sub><br>mol<sup>-1</sup> photons) | CONTROL | $0.014 \pm 0.002$ c | $0.025 \pm 0.002$ b | $0.028 \pm 0.001$ ab | $0.030 \pm 0.001$ ab | $0.031 \pm 0.003$ ab | $0.036 \pm 0.002$ a |
| | SHADE | $0.012 \pm 0.001$ a | $0.010 \pm 0.002$ a | $0.015 \pm 0.001$ a | $0.013 \pm 0.001$ a | $0.013 \pm 0.001$ a | $0.012 \pm 0.001$ a |
| | *Shade/Control* | $0.92 \pm 0.13$ a | $0.4 \pm 0.09$ b | $0.55 \pm 0.03$ b | $0.44 \pm 0.02$ b | $0.42 \pm 0.06$ b | $0.34 \pm 0.04$ b |

### Using electrochromic signal to determine the photosystem stoichiometry

Having determined the fraction (*r*) of P700 in PSI remaining reduced during illumination with moderately strong far-red light, and available for charge separation, we obtained the xenon flash-induced EC signal in the absence or presence of the far-red light (Figure S1). The amplitude of the fast rise decreased in the presence of moderately strong far-red light, being now contributed to by all the PSII and the small fraction *r* of the PSI reaction centres (Figure S2). The ratio of the PSII/PSI reaction centres was then calculated according to Equation 3 (Figure 1C; Table 2). The leaf PSII/PSI ratio was highest in *P. bisulcatum* (C_3_) relative to other control species. Shade significantly lowered leaf PSII/PSI ratio of most species except for *P. miliaceum* (NAD-ME) and *Z. mays* (NADP-ME), implying the activity of PSII and PSI were equally down-regulated under shade in those two species (Figure 1C; Tables 1 and 2). Accordingly, PSI content (obtained by dividing PSII content by the PSII/PSI ratio) declined only in those two species (*P. miliaceum* and *Z. mays*) under shade (Table 2). Otherwise, PSI did not significantly vary with photosynthetic types/subtype or light treatment (Table 2).

### Leaf electron fluxes from custom-built equipment

Leaf LEF_O2_, measured using a gas-phase O_2_ electrode, and total electron flux through PSI (ETR1) both measured at saturating irradiance varied amongst the grass species independently of the photosynthetic type (Figures 2A-B; Tables 1 and 2). In control plants, LEF_O2_ and ETR1 were highest in the two NADP-ME grasses and lowest in *M. maximus* (PCK) relative to the other species (Tables 1 and 2). Shade substantially reduced LEF_O2_ (−40% to −60%) and ETR1 (−55% to −80%) in all species, but both parameters remained higher in *Z. mays* relative to other shaded species (Figures 2A-B; Tables 1 and 2).

**Figure 2.**
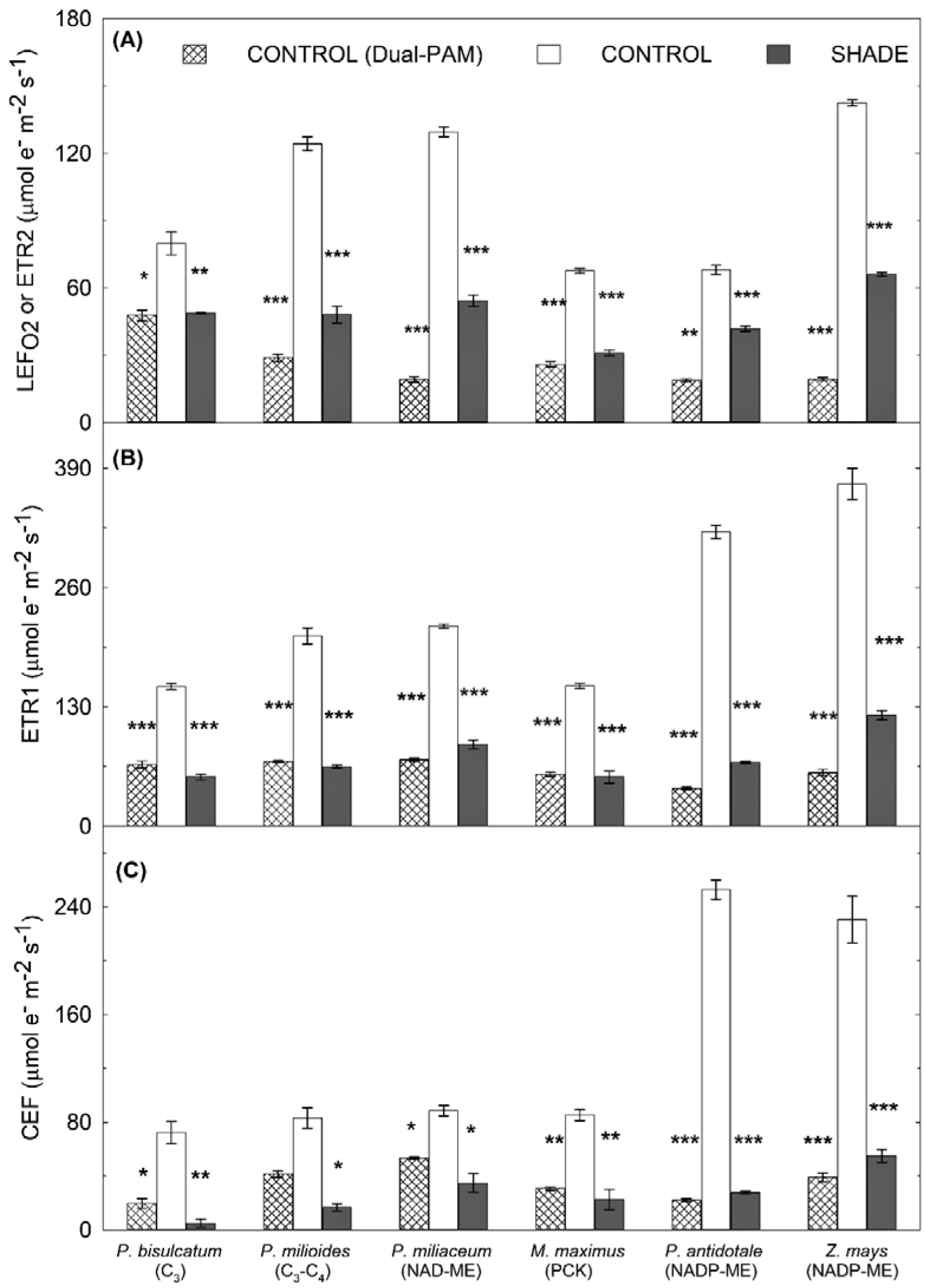
Electron fluxes in leaves of control and shade-grown C_3_, C_3_-C_4_ and C_4_ grasses. Measurements of **(A)** electron flux through both photosystems (LEF_O2_) and ETR2; **(B)** electron flux through PSI (ETR1); and **(C)** cyclic electron flux around PSI (CEF) at saturating light (2000 μmol photons m^−2^ s^−1^) in leaves of control and shade-grown C_3_, C_3_-C_4_ and C_4_ grasses. Electron fluxes (clear and grey columns) were measured using custom-built equipment for oxygen evolution (LEF_O2_) and redox kinetics of P700 (ETR1). For comparison, electron fluxes of control plants were also measured using Dual-PAM (checked columns). Each column represents the mean ± s.e. of each species (*n* = 4 plants). Statistical significance levels (t-test) for the growth condition within each species are shown and they are: * ≡*p*<0.05; ** ≡*p* < 0.01; *** ≡*p* < 0.001.

In control plants, cyclic electron flow (CEF) was highest in the two NADP-ME species, and similar amongst the other species (Figure 2C; Tables 1 and 2). Shade significantly reduced CEF in all species; the reduction was most pronounced in the C_3_ (−94%) and least in the NAD-ME species (−60%) relative to the other species, with the C_4_ species generally maintaining higher CEF under shade relative to the two non-C_4_ species (Figure 2C; Tables 1 and 2). CEF constituted a higher fraction of total electron flow (LEF_O2_ or ETR1) in NADP-ME and PCK relative to the other species, particularly under HL conditions; this fraction declined under shade except in the NAD-ME species (Tables 1 and 2). Overall, there were good linear relationships between *A* measured at HL and CEF or LEF_O2_ for individual species: however, there was no strong common relationship across the various species even within C_4_ plants (Figures S4A-B).

### Custom-built unit versus commercial equipment for measuring leaf electron fluxes

Electron flux data from leaves measured using custom-built equipment were compared with data gathered using the commercial Dual-PAM (Figures 2A-C; Tables 1, 2 and S1). In control plants, values of ETR2 from Dual-PAM were significantly lower compared to LEF_O2_ (Figures 2A and S4C; Tables 1, 2 and S1). There was also a discrepancy between ETR1 from the custom-built unit compared to ETR1 from Dual PAM (Figures 2B and S4D; Tables 1, 2 and S1). Both relationships markedly deviated from 1:1, and showed distinct relationships for certain species such as *Z. mays* and *P. milioides* (Figures S4C-D). Accordingly, CEF rates of control species were severely under-estimated by using Dual-PAM relative to the custom-built equipment (Figure 2C). For the remainder of the Results and Discussion sections we only consider data collected using custom-built equipment.

### Leaf optical properties

Absorptance spectra for all grass species were typical of vascular plants (Björkman & Demmig, 1987; Knapp & Carter, 1998). The highest total leaf-specific absorptance (up to 80% of incident irradiance) was observed in the blue (400–500 nm) and red (600–680) regions around the chlorophyll *a* peak of 660–670 nm (Figures 3 and S5). The lowest total leaf-specific absorptance (as low as 50% of incident irradiance) occurred in the green wavelengths (500-580 nm), the spectral region with the highest transmittance and reflectance (Figures 3 and S5). Shade-reduced leaf absorptance of all species in the visible region of the spectrum (400-700 nm). This reduction was more pronounced in the green region (−3% to −18%) and around the chl *a* peak (660-670 nm) with 0% to −11% reduction, in comparison to the blue region (+2% to −8%) (Figure 3). Among the shade-grown species, *P. bisulcatum* (Figure 3A; Table 1) and *P. antidotale* (Figure 3E; Table 1) had the least reductions in total absorptance (−2% and −10%, respectively), while and *P. miliaceum* had the greatest reduction (−37%) (Figure 3C; Table 1). The remaining grasses showed reductions of −20% (Figure 3D), −25% (Figure 3F) and −26% (Figure 3B) in total absorptance.

**Figure 3.**
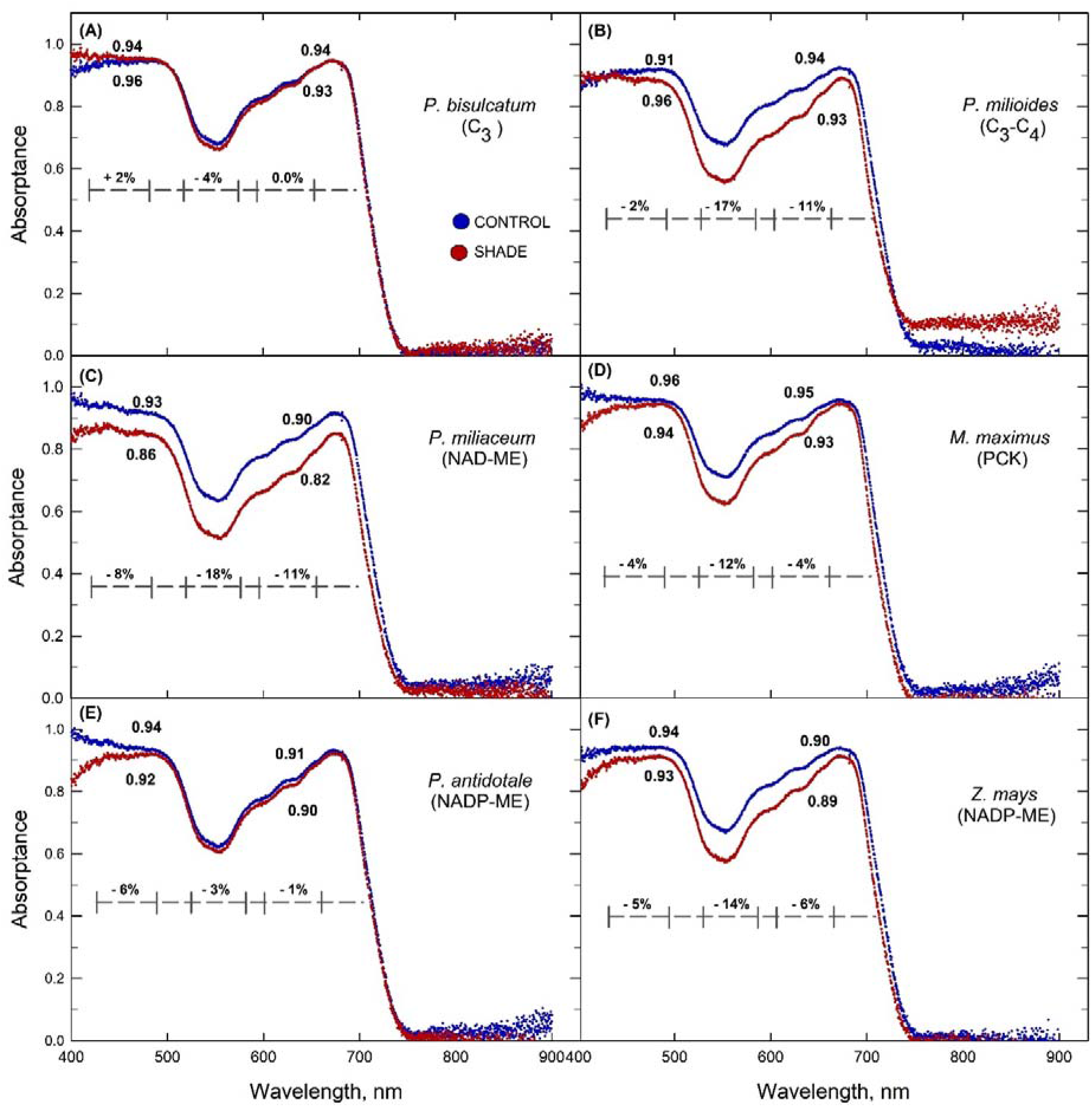
Leaf average spectral absorptance. Average leaf spectral absorptance showing the percentage decrease in absorptance in the blue (430-453 nm), green (500-580 nm), and red (640-662 nm) regions for control (blue traces) and shade- (red traces) grown (**A**) *Panicum bisulcatum* (C_3_); (**B**) *Panicum milioides* (C_3_-C_4_); (**C**) *Panicum miliaceum* (NAD-ME); (**D**) *Megathyrsus maximus* (PCK); (**E**) *Panicum antidotale* (NADP-ME); and (**F**) *Zea mays* (NADP-ME) (*n* = 4 plants). The numbers represent the specific absorptance for control and shaded leaves at 470 nm and 665 nm, corresponding to the blue and red LED light source used by the LICOR equipment during gas exchange.

### Leaf pigment content and composition

Lutein was the predominant carotenoid in all samples; the other carotenoids that accumulated, in decreasing order of abundance, were β-carotene, violaxanthin, neoxanthin, antheraxanthin and zeaxanthin (Tables 1 and 3). All species contained the same pigments, but the ratios of the individual pigments (expressed as a percentage of total carotenoid) differed between species (Tables 1 and 3). Shade reduced total leaf carotenoid and chlorophyll contents of *P. milioides, P. miliaceum* and *P. antidotale,* while chlorophyll content increased in *Z. mays* (Figures 4A-B; Table 1). The greatest decrease was observed in *P. miliaceum* with 45% and 38% decrease in leaf carotenoid and chlorophyll content, respectively (Figures 4A-B; Table 1). Shade also reduced the ratios of chlorophyll *a/b* and carotenoids/chlorophyll in all species; i.e. carotenoids became less abundant relative to chlorophylls under shade (Figure 4D; Table 1). These results are generally consistent with the literature whereby shade or low light plants have lower chlorophyll *a/b* and carotenoids/chlorophyll ratios relative to sun or high light plants (Anderson, 1986). Overall, there was a strong linear relationship between total chlorophyll and total carotenoid contents in the leaves of all species under both light treatments (Figure S6).

**Figure 4.**
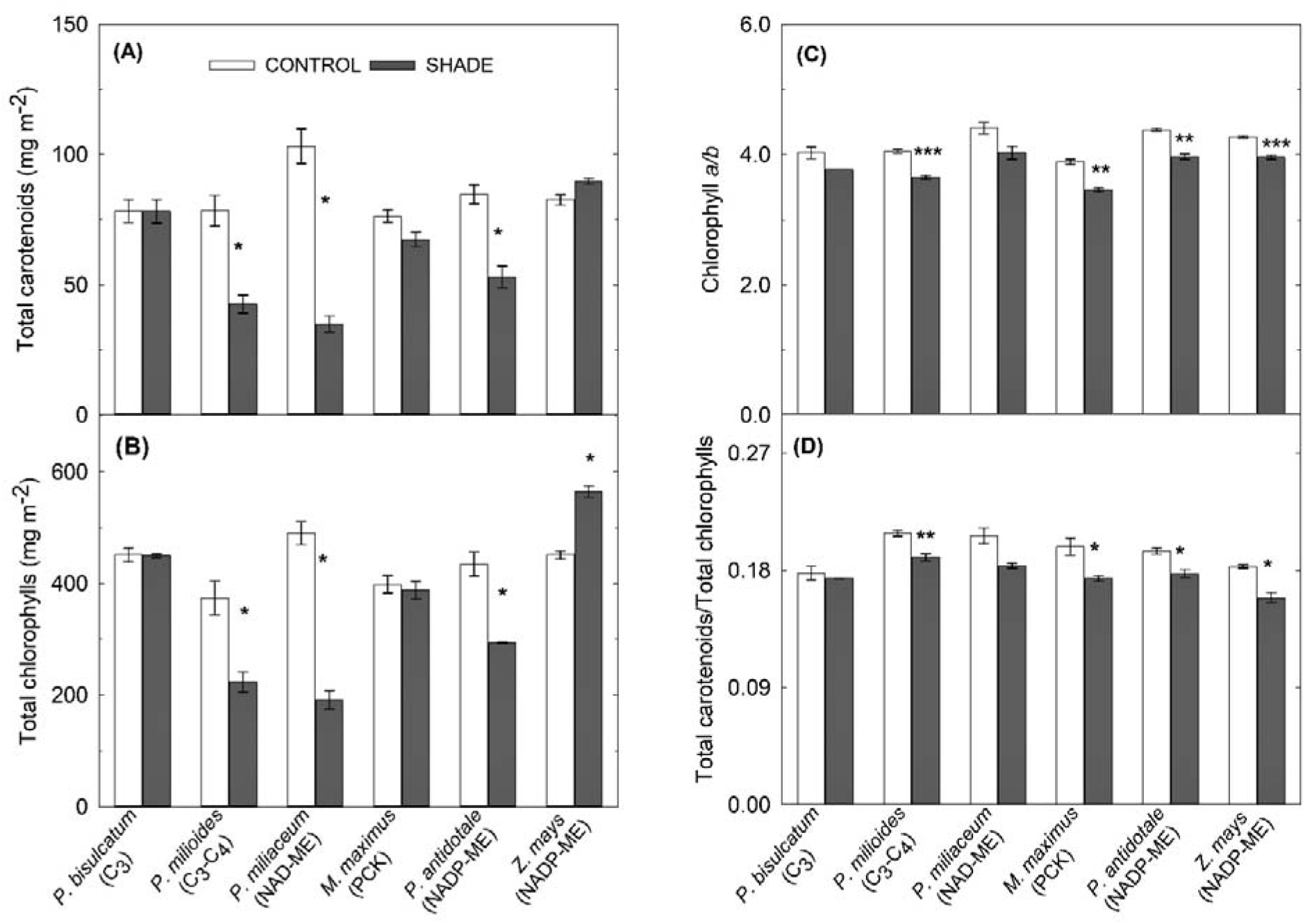
Leaf content of chlorophyll and carotenoids. Leaf **(A)** total carotenoids; **(B)** total chlorophylls; **(C)** chlorophyll *a/b* ratio; and **(D)** total carotenoid/total chlorophyll ratio in control and shade-grown C_3_, C_3_-C_4_ and C_4_ grasses. Each column represents the mean ± s.e. of each species (*n* = 4 plants). Statistical significance levels (t-test) for the growth condition within each species are shown and they are: * ≡ *p*<0.05; ** ≡ *p* < 0.01; *** ≡ *p* < 0.001.

**Table 3.** Leaf pigments in representative C_3_, C_3_-C_4_, and C_4_ grasses. Leaf pigments in representative C_3_, C_3_-C_4_, and C_4_ species grown under control (~800 μmol photons m^−2^ s^−1^) or shaded (~200 μmol photons m^−2^ s^−1^) conditions. Different letters indicate significantly different (*p*<0.05) ranking within each species using multiple-comparison Tukey’s post-hoc test. Values are means ± s.e. of species (*n* = 4 plants).

| Parameter | Light | *P. bisulcatum* | *P. milioides* | *P. miliaceum* | *M. maximus* | *P. antidotale* | *Z. mays* |
| --- | --- | --- | --- | --- | --- | --- | --- |
|  |  | C<sub>3</sub> | C<sub>3</sub>-C<sub>4</sub> | NAD-ME | PCK | NADP-ME | NADP-ME |
| Neoxanthin (%) | CONTROL | 9.5 ± 0.2 ab | 10 ± 0.2 a | 8.4 ± 0.2 c | 9.9 ± 0.3 a | 8.6 ± 0.2 bc | 8.0 ± 0.1 c |
| | SHADE | 10.3 ± 0.0 a | 10.6 ± 0.3 a | 8.3 ± 0.2 b | 10.7 ± 0.3 a | 8.6 ± 0.4 b | 8.1 ± 0.3 |
| | Shade/Control | 1.09 ± 0.03 a | 1.06 ± 0.04 a | 1 ± 0.05 a | 1.09 ± 0.06 a | 1 ± 0.07 a | 1.01 ± 0.03 a |
| Violaxanthin (%) | CONTROL | 20.5 ± 0.5 a | 20.3 ± 0.8 ab | 19.5 ± 0.6 ab | 18.5 ± 0.5 ab | 18.5 ± 0.5 ab | 17.1 ± 0.5 b |
| | SHADE | 15.9 ± 0.0 b | 17.9 ± 0.3 a | 15.9 ± 0.2 b | 9.2 ± 0.2 e | 14.0 ± 0.4 c | 12.0 ± 0.2 d |
| | Shade/Control | 0.78 ± 0.02 ab | 0.89 ± 0.05 a | 0.82 ± 0.03 ab | 0.5 ± 0.01 c | 0.76 ± 0.04 ab | 0.7 ± 0.02 b |
| Antheraxanthin (%) | CONTROL | 1.7 ± 0.0 ab | 1.6 ± 0.0 ab | 1.8 ± 0.1 ab | 1.6 ± 0.1 b | 1.7 ± 0.0 ab | 1.8 ± 0.0 a |
| | SHADE | 1.5 ± 0.0 a | 1.6 ± 0.0 a | 2.0 ± 0.0 a | 1.8 ± 0.3 a | 1.9 ± 0.1 a | 1.7 ± 0.0 a |
| | Shade/Control | 0.87 ± 0.01 a | 1.01 ± 0.03 a | 1.11 ± 0.04 a | 1.17 ± 0.22 a | 1.11 ± 0.07 a | 0.97 ± 0.03 a |
| Lutein (%) | CONTROL | 36.8 ± 0.7 c | 37.3 ± 0.3 c | 39.2 ± 0.8 bc | 40.9 ± 0.9 ab | 39.5 ± 0.4 abc | 42.1 ± 0.4 a |
| | SHADE | 42.4 ± 0.0 c | 43.0 ± 0.2 c | 43.7 ± 0.4 bc | 47.8 ± 0.5 a | 45.1 ± 0.4 b | 48.0 ± 0.2 a |
| | Shade/Control | 1.15 ± 0.02 a | 1.15 ± 0.01 a | 1.12 ± 0.02 a | 1.17 ± 0.04 a | 1.14 ± 0.02 a | 1.14 ± 0.01 a |
| Zeaxanthin (%) | CONTROL | 1.5 ± 0.1 b | 3.1 ± 0.3 a | 1.3 ± 0.3 b | 0.9 ± 0.1 b | 1.4 ± 0.4 b | 1.6 ± 0.2 b |
| | SHADE | 1.7 ± 0 b | 0.6 ± 0.1 c | 0.3 ± 0.0 c | 4.5 ± 0.5 a | 0.6 ± 0.1 c | 0.6 ± 0.1 c |
| | Shade/Control | 1.16 ± 0.07 b | 0.22 ± 0.05 b | 0.28 ± 0.06 b | 5.42 ± 0.86 a | 0.68 ± 0.31 b | 0.42 ± 0.07 b |
| β-Carotene (%) | CONTROL | 30 ± 0.2 ab | 27.8 ± 0.2 b | 29.9 ± 0.5 ab | 28.3 ± 1.0 ab | 30.3 ± 0.5 a | 29.5 ± 0.4 ab |
| | SHADE | 28.2 ± 0.0 b | 26.3 ± 0.2 c | 29.8 ± 0.5 a | 25.9 ± 0.2 c | 29.8 ± 0.4 a | 29.6 ± 0.2 a |
| | Shade/Control | 0.94 ± 0.01 a | 0.95 ± 0.01 a | 1 ± 0.02 a | 0.92 ± 0.04 a | 0.98 ± 0.03 a | 1 ± 0.01 a |
| chlorophyll a<br>(mg m<sup>-2</sup>) | CONTROL | 382.9 ± 21.8 a | 332.2 ± 36.2 a | 366 ± 35 a | 284.5 ± 33.7 a | 378.1 ± 27.4 a | 365.9 ± 6.0 a |
| | SHADE | 355.6 ± 2.5 ab | 175.8 ± 14.0 d | 179.8 ± 28.4 d | 301 ± 11.8 bc | 212.6 ± 21.9 cd | 450.7 ± 8.3 a |
| | Shade/Control | 0.94 ± 0.05 ab | 0.55 ± 0.07 c | 0.53 ± 0.15 c | 0.75 ± 0.03 bc | 0.81 ± 0.07 bc | 1.23 ± 0.01 a |
| chlorophyll b<br>(mg m<sup>-2</sup>) | CONTROL | 95.2 ± 6.4 a | 82.3 ± 9.5 a | 83.4 ± 8.7 a | 73.3 ± 8.8 a | 86.27 ± 5.95 a | 85.6 ± 1.3 a |
| | SHADE | 94.3 ± 0.6 ab | 48.2 ± 4.0 c | 44.4 ± 6.5 c | 87 ± 3.7 b | 53.74 ± 5.92 c | 113.9 ± 1.6 a |
| | Shade/Control | 1 ± 0.07 abc | 0.61 ± 0.08 cd | 0.58 ± 0.17 d | 0.74 ± 0.02 bcd | 1.02 ± 0.09 ab | 1.33 ± 0.01 a |

### Activity of photosynthetic enzymes

Among all control species, *P. milioides* had the highest leaf Rubisco content (37.4 ± 5.6 μmol sites m^−2^) and Rubisco activity. Among the control C_4_ species, *P. miliaceum* had the highest Rubisco content (8.8 ± 0.5 μmol sites m^−2^) while *Z. mays* had the highest Rubisco activity (Tables 1 and 4). In control C_4_ species, PEPC activity was highest in *Z. mays* and lowest in *P. miliaceum* (Tables 1 and 4). Overall, shade reduced soluble protein content and photosynthetic enzyme activity to a greater extent in C_3_, C_3_-C_4_ and C_4_-NAD-ME relative to the other C_4_ species (Tables 1 and 4). In particular, Rubisco activity decreased by 73-80% in *P. bisulcatum, P. milioides* and *P. miliaceum,* and by 44-66% in the PCK and NADP-ME C_4_ grasses (Tables 1 and 4). Shade reduced PEPC activity by 62–76% in the C_4_ species (Tables 1 and 4).

**Table 4.**
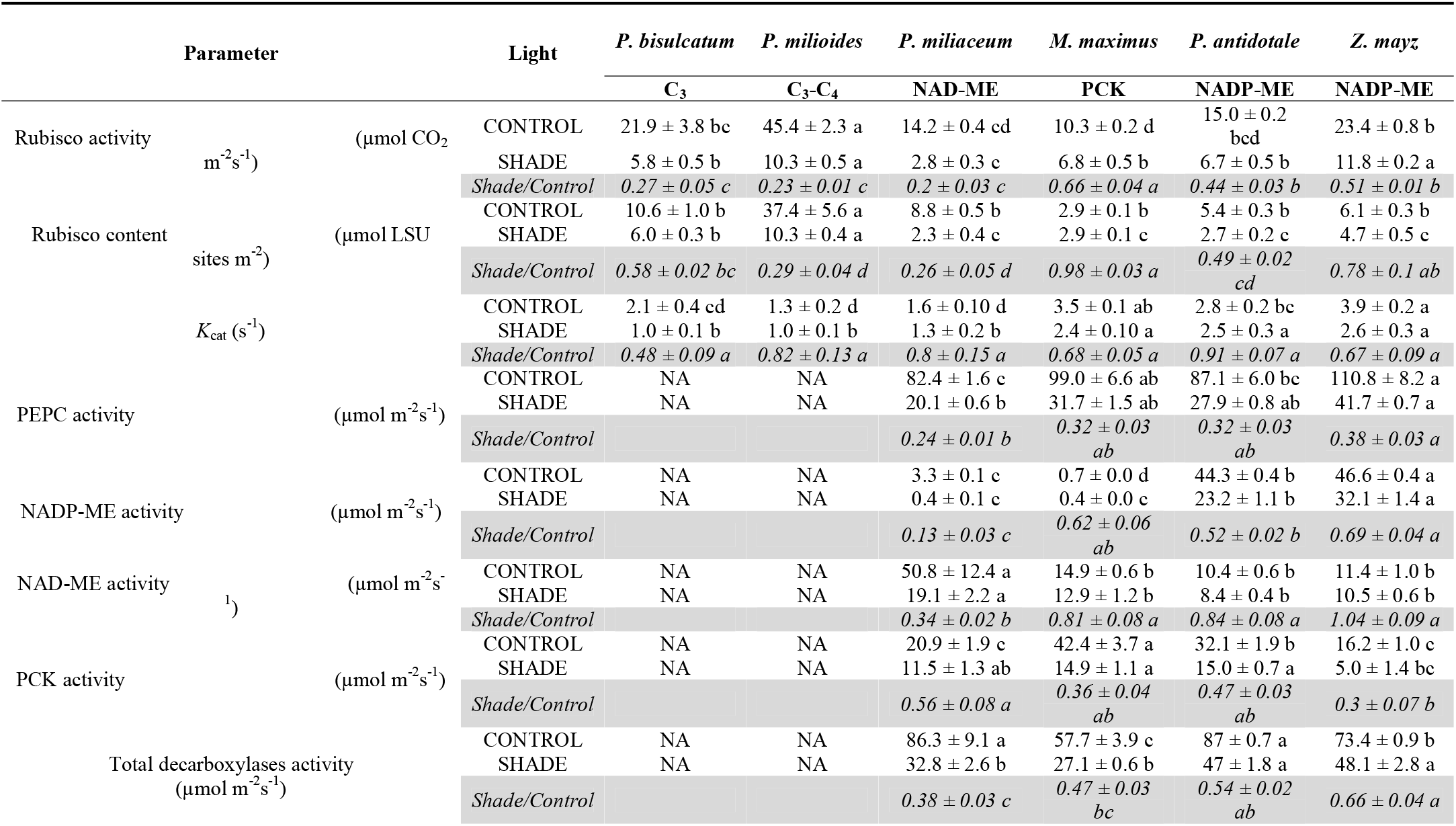
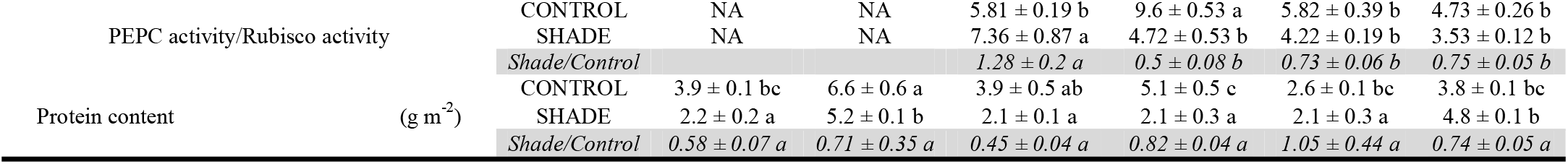
Activities of Rubisco, PEPC and decarboxylases and soluble protein content. Activities of Rubisco, PEPC and decarboxylases and soluble protein content measured in leaves of representative C_3_, C_3_-C_4_, and C_4_ grasses grown under control (~800 μmol photons m^−2^ s^−1^) or shaded (~200 μmol photons m^−2^ s^−1^) conditions. Different letters indicate significantly different (*p*<0.05) ranking within each species using multiple-comparison Tukey’s post-hoc test. Values are means ± s.e. of species (*n* = 4 plants; NA = not applicable).

Activities of NADP-ME, PCK, and NAD-ME enzymes were dominant in the species with the respective subtype. In addition, substantial PCK activity (16–32 μmol m^−2^ s^−1^) and NAD-ME activity (42–58 μmol m^−2^ s^−1^) operated as secondary decarboxylases in all control C_4_ species (Figure 5B; Tables 1 and 4). Shade reduced the total decarboxylase (−36% to −59%) and specific decarboxylase activity of NADP-ME (−31% to −48%) less than PCK (−65%) and NAD-ME (−62%) species (Figure 5A; Tables 1 and 4). Shade generally reduced the contribution of the secondary decarboxylases in the NAD-ME and PCK, but not NADP-ME, species (Figure 5B; Tables 1 and 4).

**Figure 5.**
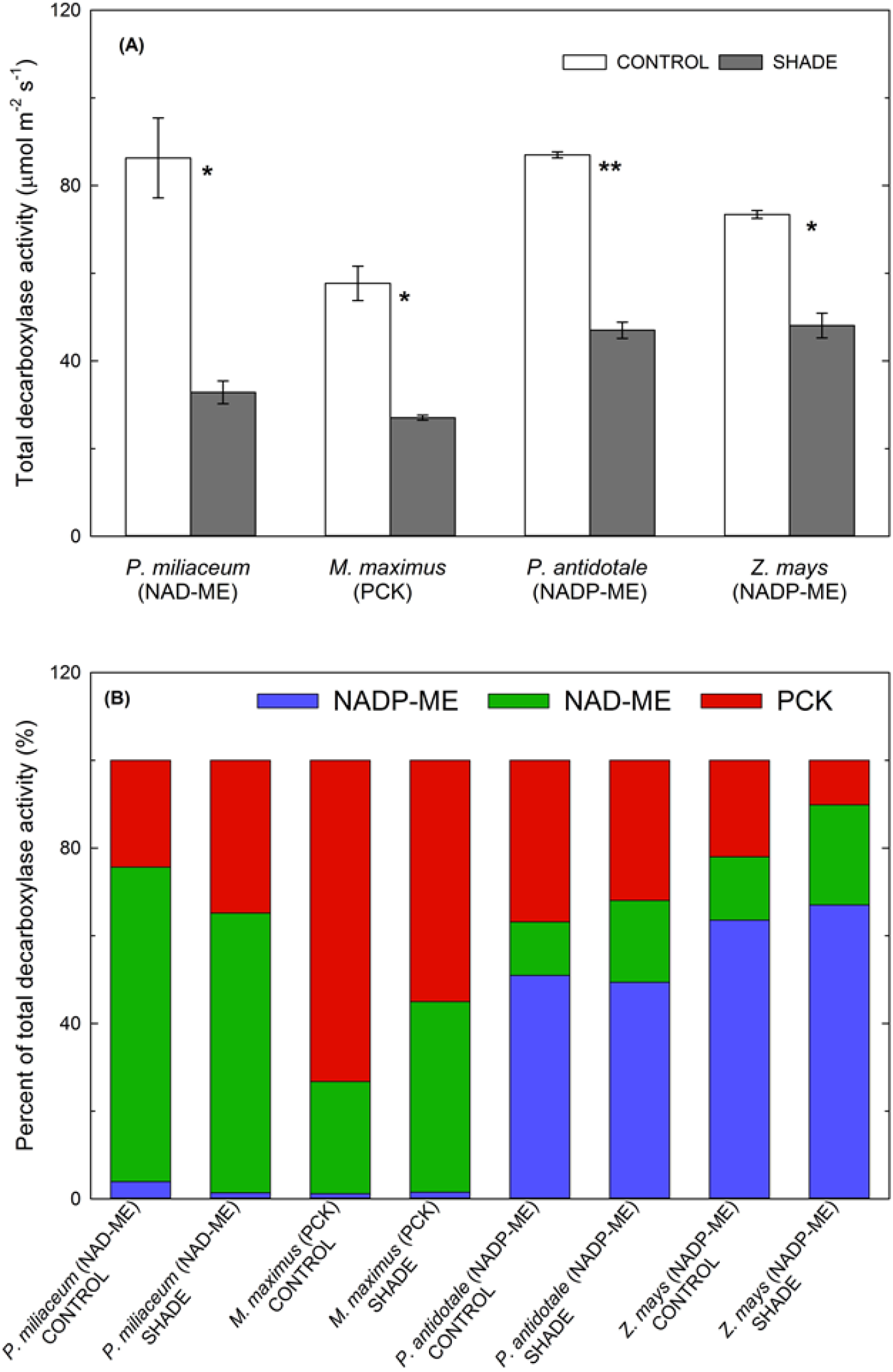
Activities of the C_4_ decarboxylases. Activity of the C_4_ decarboxylases **(A)** and percentage contribution of each C_4_ decarboxylase to the total decarboxylase activity **(B)** in control and shade-grown C_4_ grasses. Each column represents the mean ± s.e. of each species (*n* = 4 plants). Statistical significance levels (t-test) for the growth condition within each species are shown and they are: * ≡*p*<0.05; ** ≡*p* < 0.01; *** ≡*p* < 0.001.

### Quantum yield and assimilation quotient

Quantum yield (*QY*) was calculated as the ratio between CO_2_ assimilation rates measured at ambient CO_2_ and high (*A*1000) or growth irradiance (*A*growth) to absorbed irradiance (Figure 1D; Tables 1 and 2). In control plants, *QY*1000 and *QY*growth were generally lowest in C_3_, intermediate in C_3_-C_4_ and highest in C_4_ species. Under shade, *QY*1000 and *QY*growth were lower in C_3_ relative to C_4_ species (Figure 1D; Tables 1 and 2). Within the C_4_ species, there was a non-significant trend for *QY* to be highest in NADP-ME, intermediate in PCK and lowest in NAD-ME species (Figure 1D; Tables 1 and 2), in line with previous observations (Ehleringer & Pearcy, 1983; Sonawane *et al.*, 2018). The assimilation quotient (*AQ*_1000_) was calculated as the ratio of *A*_1000_ to LEF_O2_/4. *AQ*_1000_ was highest in the PCK and NADP-ME species (although lower in *Z. mays* than *P. antidotale*) relative to the NAD-ME and the two non-C_4_ species (Tables 1 and 2). Lower *AQ*_1000_ in *Z. mays* relative to the other NADP-ME species may reflect the high photosynthetic capacity of maize which would normally require above ambient CO_2_ for saturation. In turn, lower *AQ*1000 and *QY*1000 in C_3_ and C_3_-C_4_ species reflect their high photorespiration rates at ambient CO_2_ relative to C_4_ species.

## DISCUSSION

This study investigated whether variations in light conversion efficiency and susceptibility to low light depend on photosynthetic types or subtypes. Accordingly, we grew six related C_3_, C_3_-C_4_ and C_4_ Paniceae grasses, including representatives of all three C_4_ subtypes, under control (full sunlight) and shade (20% sunlight) conditions and measured leaf chlorophyll fluorescence, photosynthesis, electron transport, absorptance and pigment contents.

### The C_3_ and C_3_-C_4_ species showed distinct photosynthetic responses to shade

Under control (high light) conditions, the C_3_-C_4_ species had intermediate photosynthetic rates, *QY* and *AQ* relative to C_3_ and C_4_ species (Figure 1, Table 2), reflecting reduced photorespiration rates due to the operation of the glycine shuttle in C_3_-C_4_ plants and the CCM in C_4_ plants (Sage *et al.*, 2012). The advantage of the C_3_-C_4_ over C_3_ species disappeared under shade due to greater sensitivity of C_3_-C_4_ photosynthesis (Figure 1A). In addition, C_3_-C_4_ plants required higher CEF under shade, possibly to cover the costs of the C_3_-C_4_ shuttle and the operation of photosynthetic reactions in two cell types relative to C_3_ photosynthesis (Pinto *et al.*, 2011; Sage *et al.*, 2012). Relative to C_4_, the C_3_ and C_3_-C_4_ species had higher leaf PSII contents, partially reflecting the higher Rubisco contents in these species, as well as the importance of PSII in balancing energy in these plants through photochemical and non-photochemical quenching (Ruban *et al.*, 2012). Shade significantly lowered PSII/PSI ratio in both C_3_ and C_3_-C_4_ species such that calculated PSI content was higher in the C_3_-C_4_ than C_3_ species but unaltered by shade (Figure 1C; Table 2). Greater PSI activity in the C_3_-C_4_ species may be required to support more CEF and higher ratio of ATP to NADPH particularly under shade. However, for most species the decrease in PSII/PSI ratio was explained by fewer PSII units (Figure 1B) with a larger light-harvesting antenna, relative to PSI which was little affected (Anderson *et al.*, 1988; Chow *et al.*, 1990; Hihara & Sonoike, 2001; Melis, 1991; Leong & Anderson 1984; Chow *et al.* 1988).

### The two NADP-ME species had higher CEF, AQ and QY relative to other C_4_ species

Previous studies demonstrated that low light compromised the photosynthetic efficiency most in NAD-ME followed by PCK, but not in NADP-ME grasses (Sonawane *et al.*, 2018). The NAD-ME species showed increased BS CO_2_ leakiness, greater reduction in *QY* and greater increase in carbon isotope discrimination under shade, which were interpreted as lower CCM efficiency and imbalance between the C_3_ and C_4_ cycles (Bellasio & Griffiths, 2014a, 2014b; Sonawane *et al.*, 2018). Here, we show that higher efficiency of NADP-ME (and conversely, lower efficiency of NAD-ME) species under shade was also contributed by the light energy absorption and conversion characteristics of the C_4_ subtypes.

The NADP-ME subtype is characterised by low PSII content in BS chloroplasts (Anderson *et al.*, 1971; Dengler *et al.*, 1994; Drozak & Romanowska, 2006; Ghannoum *et al.*, 2005; Hernández-Prieto *et al.*, 2019; Romanowska *et al.*, 2008). As such, it has long been hypothesised that NADP-ME species require higher PSI efficiency (Furbank *et al.*, 1990). This was reflected in the higher rates of ETR1 and CEF under saturating light in the two NADP-ME species compared to other species (Figure 2B; Table 2). These observations support the findings that BS chloroplasts of NADP-ME species have high PSI content (Ku *et al.*, 1974; Romanowska *et al.*, 2006, 2008; Schuster *et al.*, 1985; Woo *et al.*, 1970), and hence are expected to generate ATP through CEF. Operation of enhanced CEF around PSI might partially explain the higher *QY* and *AQ* in NADP-ME species. (Yin & Struik, 2018) suggested that extra ATP required in C_4_ photosynthesis for the CCM comes from CEF, and that ~50% of electron flux in NADP-ME species is CEF which predominantly occurs in BSCs. Indeed, Table 2 shows that in control growth light, the two NADP-ME species had CEF/ETR1 ratios of 79% (*P. antidotale*) and 62% (*Zea mays*), and which were higher than in the other C_4_ (41-56%) and non-C_4_ (40-48%) species. Shade reduced ETR1 and CEF the greatest in the two NADP-ME species, with the CEF/ETR1 ratio falling to about 40-45% among the C_4_ species) (Figure 2; Table 2), highlighting the dependence of photosynthetic rates on CEF in this subtype. Accordingly, we observed a strong correlation between CEF and photosynthetic rate of C_4_ species under high irradiance (Figure S4A). It is worth noting that on a total leaf level, enhanced CEF in the two NADP-ME species was not accompanied with greater PSI content, implying greater PSI efficiency in this subtype (Table 2).

In maize (NADP-ME), (Bellasio & Griffiths, 2014c) suggested that the operation of two decarboxylase systems (NADP-ME and PCK) in the BSC increases the flexibility of energy, particularly NADPH, supply. The two NADP-ME species had significant PCK and NAD-ME activities, but no appreciable changes in the decarboxylase contribution between control and shade treatments was observed in any of the C_4_ species (Figure 5). Nevertheless, employing mixed C_4_ pathways, which include the PCK type, theoretically decreases the need to maintain high concentration gradients of transport metabolites and affords high photosynthetic efficiency under a broad range of light regimes (Wang *et al.*, 2014). This may partially explain the high energy efficiency in NADP-ME and PCK relative to NAD-ME species (Table 2). Higher *AQ* and *QY* support the view that these NADP-ME and PCK subtypes have reduced leakiness (Sonawane *et al.*, 2018), even though differences in leakiness amongst the C_4_ subtypes have been difficult to detect using stable isotopic techniques (Cousins *et al.*, 2008; Henderson *et al.*, 1992).

### Among C_4_, the NAD-ME species had lower energy efficiency and greater sensitivity to shade

Whilst leaf PSII/PSI ratio was similar among the C_4_ grasses, PSII content was lowest and most reduced by shade in the NAD-ME species. This may seem surprising given that PSII activity is generally highest in isolated M and BS chloroplasts of PCK *M. maximus* and NAD-ME *P. miliaceum* than NADP-ME *Z. mays* (Drozak & Romanowska, 2006). CEF was least reduced in *P. miliaceum* while PSI content was most reduced by shade among the C_4_ species (Table 2). This seems puzzling; however, it is not clear what limits CEF. It is possible that the redox poise of components in the cycle is more limiting than PSI content (Allen, 2003).

Reduced leaf Y(II), *F*_v_/*F*_m_, LEF_O2_ and PSII content (Table 2) under shade reflects the greater photosynthetic inhibition of NAD-ME species to low light relative to the other C_4_ subtypes (Sonawane *et al.*, 2018), and might also be an acclimation response to prevent photoinhibition (Park *et al.*, 2016). The greater shade sensitivity of *P. miliaceum* may be attributable to the inefficiency of the CCM or light harvesting components of NAD-ME subtype, and its inability to adjust under shade (limited plasticity) since it is more adapted to open habitats relative to the other two C_4_ subtypes (Liu *et al.*, 2012; Schulze *et al.*, 1996; Vogel *et al.*, 1986). It also has two fully fledged linear electron transport systems in both MC and BSC which both need to adjust under shade. This is not the case for the NADP-ME subtype, and somewhat intermediate for the PCK subtype (Ghannoum *et al.*, 2005; Pinto *et al.*, 2011). *P. miliaceum* also showed the greatest decrease in the activity of Rubisco and decarboxylases, similar to what has previously been reported (Sonawane *et al.*, 2018). In addition, the NAD-ME species had lower *AQ* than PCK and NADP-ME species and lower *QY* than NADP-ME species, all indicative of the greater sensitivity of NAD-ME species to shade (Sonawane *et al.*, 2018).

### Changes in leaf pigments under shade reflect increased light harvesting rather than photoprotection

Photosynthetic pigments play an important role in *QY* variations because they balance the absorption of light energy. Strong correlations between total chlorophyll and total carotenoid contents across species and treatments (Figure S6) suggest the related functions of these pigments in leaves. Lutein was the most abundant leaf carotenoid (Table 3), as it often accounts for >50% of the total carotenoid pool in plants. Lutein is localized in LHCs and is the only carotenoid detected in the PSII core (Bassi *et al.*, 1993; Kühlbrandt *et al.*, 1994). The greater abundance of lutein in C_4_ compared to C_3_ and C_3_-C_4_ leaves suggests that leaves of high light adapted C_4_ plants (Edwards *et al.*, 2010; Ehleringer *et al.*, 1997) have more LCHII and are more efficient in protecting PSII cores against photoinhibition.

Violaxanthin, another abundant pigment detected in the leaf of all species (Table 3), plays an important role in the thermal dissipation of excessive energy (Jin, 2003; Yamamoto, 1979) via the xanthophyll cycle (Demmig-Adams, 1998; Demmig & Bjorkman, 1987). Zeaxanthin is generally not formed in leaves at low irradiance, but as the amount of energy captured begins to exceed that which can be used in photosynthesis, more zeaxanthin is formed from violaxanthin (Demmig-Adams & Adams, 1992). Hence, low amounts of zeaxanthin in leaves of C_3_ and C_4_ relative to the C_3_-C_4_ species might indicate lower light stress under the growing conditions (Table 3).

Leaf chl *a/b* and carotenoids/chl ratios decreased under shade in all except the C_3_ species (Figures 4C-D). Chl *a* is the most common photosynthetic pigment and absorbs blue, red and violet wavelengths in the visible spectrum. Chl *b* primarily absorbs blue light, extending the absorption spectrum of chl *a*. Chl *a/b* ratios are greater in the core complex than the light-harvesting complex for both photosystems (Thornber, 1986), and are positively correlated with the ratio of PSII cores to light harvesting chlorophyll-protein complex (LHCII) (Hikosaka & Terashima, 1995). LHCII contains the majority of chl *b*, and consequently has a lower chl *a/b* ratio than other chlorophyll binding proteins associated with PSII (Evans, 1989; Green & Durnford, 1996). Hence, lower chl *a/b* ratios under shade imply an increased proportion of light-harvesting complexes relative to reaction centres (Green & Durnford, 1996; Hikosaka & Terashima, 1995), and this might be an adaptation to broaden the spectral range over which PSI and PSII absorb light (Yamazaki *et al.*, 2005). Reduced carotenoids/chl ratios (Figure 4D) reflects the strategy of shade plants to prioritise light harvesting (through chl) more than photoprotection (through carotenoids).

### Shade reduced leaf absorptance most in the NAD-ME and least in the C_3_ species

Shade responses of leaf absorptance and pigments were pronounced in the C_3_-C_4_ and C_4_ species, but virtually unaffected in the C_3_ species. This response represents the greater sensitivity of the C_4_ pathways to low light (Sharwood *et al.*, 2014; Sonawane *et al.*, 2018). Lower absorptance (increased leaf reflectance) under shade can partially be explained by lower chlorophyll and carotenoid contents (Figure 3). Chlorophyll content is a sensitive indicator of plant stress (Evans, 1993; Lin & Ehleringer, 1983; Vogelmann, 1993). Leaf spectral properties are more consistently altered in response to stress in the visible wavelengths than in the remainder of the incident solar spectrum (Carter, 1993; Carter *et al.*, 1992). Similarly, leaf absorptance in the current study noticeably decreased in the green region (500-580 nm), where non-photosynthetic pigments such as anthocyanins also absorb (Gould *et al.*, 1995; Paradiso *et al.*, 2011; Smillie & Hetherington, 1999), and in the chl *a* region (660-670 nm), which reflects the lower chl *a/b* ratio in shaded plants. These changes can also be due to alterations in leaf structure under shade. Reduced leaf thickness and changes in arrangement of cells within a leaf which can increase transmitted light, thus preventing photoinhibition (Vogelmann, 2003).

Shade led to a smaller reduction in leaf absorptance of the NADP-ME and PCK relative to the NAD-ME species (Figure 3). In maize, leaf pigments increased and suffered no reduced light absorptance under shade (Figures 3 and 4), indicating that shade acclimation involved optimal nitrogen allocation to chloroplast pigment-proteins in order to balance energy capture and energy transfer as previously suggested (Evans, 1989; Hikosaka & Terashima, 1995). Such adjustments are advantageous for leaves which developed under shade and, hence are more susceptible to photosystem damage than control leaves. These plants cannot dissipate excess light energy due to reduced photosynthesis, so NPQ involving carotenoids was needed. Increased chlorophyll content is also an adaptive and common response to shade, since they can provide a higher light harvesting capacity in low-light environments (Lei & Lechowicz, 1997; Lei *et al.*, 1996). The relatively greater decrease in total pigment in shaded *P. miliaceum* suggests a low capacity to acclimate under shade in comparison to other C_4_ subtypes and C_3_ and C_3_-C_4_ species (Figure 4).

### Comparison between custom-built and commercial equipment

The common approach to calculate CEF is to determine ETR2 and then subtracting it from ETR1 using chl fluorescence. However, (Fan *et al.*, 2016) pointed out that there are some uncertainties for using chl fluorescence in estimating ETR2. One of these is an underestimation of the fluorescence signal detected from the leaf because this signal comes from an unspecified (shallow) depth in the leaf tissue and that depth may vary during the course of an experiment, for example, due to chloroplast movement (Sato & Kadota, 2006). Subtracting ETR2 based on fluorescence measurement from ETR1 then gives an overestimation of CEF, leading to underestimation of CEF (Figure 2C). To avoid this, we calculated ETR2 based on whole-tissue measurement of the gross rate of oxygen evolution (LEF_O2_). According to (Fan *et al.*, 2016), this can be validly compared with ETR1 obtained from Y(I) because the P700^+^ signal is also a whole-tissue measurement, since 820 and 870 nm measuring beams are only weakly absorbed by the leaf tissue and are, therefore, multiply scattered in the tissue until they are finally absorbed; subtraction of LEF_O2_ from ETR1 is then valid, as both refer to the same leaf tissue.

Estimation of ETR1 by two different equipment used the same principle of the Y(I)-based electron flux. However, the source of actinic light and saturating pulse is different between the two pieces of equipment. Dual-PAM uses the red light as a source of actinic and saturating light while the custom-built equipment uses white light from a halogen lamp. ETR1 is directly estimated by Dual-PAM using equation (6), utilising 0.5 as a default value for *f*_I_. The custom-built unit separately measures the components of Y(I) and the *f*_I_ values were experimentally determined (Sagun *et al.*, 2019). Further, the custom-built unit uses a strong far-red pulse that preceded the saturating pulse in the measurement of 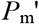, a necessary step to maximally oxidize P700 in the presence of actinic light on adding the saturating pulse (Siebke *et al.*, 1997). These factors gave different values of ETR1 which then affected CEF calculations (Figure S4D).

The ratio of PSII/PSI was independently measured using the electrochromic signal (ECS) which reflects trans-membrane charge transfer through the thylakoid membrane (Witt, 1979). The reliability of this method was evaluated by (Fan *et al.*, 2007) using spinach leaves by comparing the calculated values with two other methods, namely electron paramagnetic resonance (EPR) (Danielsson *et al.*, 2004) and by separately determining the content of functional PSII and PSI in leaf segments by the O_2_ yield per single turnover-flash and by photo-oxidation of P700 in thylakoids isolated from the same leaf. They found that the ratios obtained using ECS were comparable to the values obtained using the two other approaches. The ECS offers some advantages, being a “voltmeter reading” determined by the number of reaction centres of either photosystem present, and corresponding to a change in delocalized electric potential difference generated across the thylakoid membrane upon charge separation in the reaction centres. Further, it can also be applied to leaf segments without the need to isolate thylakoid membranes, thus avoiding any loss of PSI complexes via breakage of stromal lamellae from the membrane system, or decline of PSII activity during isolation of thylakoids.

## CONCLUSIONS

Although photochemical reactions were more efficient in C_4_ than C_3_ and C_3_-C_4_ species under sun and shade conditions, leaf pigments and absorptance were little affected in the C_3_ species. Light absorptance and conversion efficiency were generally superior in NADP-ME followed by PCK than NAD-ME species under shade. Higher *QY* and *AQ* of NADP-ME species might be associated with the comparatively greater plastic adjustments in the light harvesting components of leaves under varying environmental conditions. This important acclimation characteristic maximises light energy absorption while preventing photoinhibition under stress conditions such as long-term shade. This ensures the light reactions can still provide the right ATP/NADPH ratio even under limited supply of light energy.

## ACKNOWLEDGEMENTS

This research was funded by the Australian Research Council Centre of Excellence for Translational Photosynthesis (CE140100015) awarded to OG. JVS gratefully acknowledges the award of a Higher Degree Research Scholarship funded through the Centre of Excellence for Translational Photosynthesis and Western Sydney University.

